# A global survey of arsenic related genes in soil microbiomes

**DOI:** 10.1101/445502

**Authors:** Taylor K Dunivin, Susanna Y Yeh, Ashley Shade

**Affiliations:** Department of Microbiology and Molecular Genetics, Michigan State University, East Lansing MI 48824; Environmental and Integrative Toxicological Sciences Doctoral Program, Michigan State University, East Lansing, MI 48824, USA; Institute for Cyber-Enabled Research, Michigan State University, East Lansing, MI 48824, USA; Program in Ecology, Evolutionary Biology and Behavior, Michigan State University, East Lansing, MI 48824, USA; Department of Plant, Soil, and Microbial Sciences, Michigan State University, East Lansing, MI 48824, USA; Plant Resilience Institute, Michigan State University, East Lansing, MI, 48834, USA

**Keywords:** Arsenic, functional gene, bioinformatics, targeted gene assembly, horizontal gene transfer, biogeography, phylogeny, phylogenetic diversity, resistome, plasmid

## Abstract

Environmental resistomes include transferable microbial genes. One important resistome component is resistance to arsenic, a ubiquitous and toxic metalloid that can have negative and chronic consequences for human and animal health. The distribution of arsenic resistance and metabolism genes in the environment is not well understood. However, microbial communities and their resistomes mediate key transformations of arsenic that are expected to impact both biogeochemistry and local toxicity. We examined the phylogenetic diversity, genomic location (chromosome or plasmid), and biogeography of arsenic resistance and metabolism genes in 922 soil genomes and 38 metagenomes. To do so, we developed a bioinformatic toolkit that includes BLAST databases, hidden Markov models and resources for gene-targeted assembly of nine arsenic resistance and metabolism genes: *acr3, aioA, arsB, arsC* (grx), *arsC* (trx), *arsD, arsM, arrA*, and *arxA*. Though arsenic related genes were common, they were not universally detected, contradicting the common conjecture that all organisms have them. From major clades of arsenic related genes, we inferred their potential for horizontal and vertical transfer. Different types and proportions of genes were detected across soils, suggesting microbial community composition will, in part, determine local arsenic toxicity and biogeochemistry. While arsenic related genes were globally distributed, particular sequence variants were highly endemic (e.g., *acr3*), suggesting dispersal limitation. The gene encoding arsenic methylase *arsM* was unexpectedly abundant in soil metagenomes (median 48%), suggesting that it plays a prominent role in global arsenic biogeochemistry. Our analysis advances understanding of arsenic resistance, metabolism, and biogeochemistry, and our approach provides a roadmap for the ecological investigation of environmental resistomes.

## Background

Microbial communities drive global biogeochemical cycles through diverse functions. The biogeography of functional genes can help to predict and manage the influence of microbial communities on biogeochemical cycling [1]. These trait-based analyses require that the functional genes are well-characterized from both evolutionary and genetic perspectives [2]. The arsenic resistance and metabolism genes exemplify a suite of well-characterized functional genes that have consequences for biogeochemistry. Arsenic is a toxic metalloid that, upon exposure, can have negative effects for all life, including humans, livestock, and microorganisms. The toxicity and mobility of arsenic depends, in part, on its oxidation state: the trivalent arsenite is more mobile and more toxic than the pentavalent arsenate [3]. The toxicity of methylated arsenic species varies with oxidation state and number of methyl groups (monomethyl, dimethyl, trimethyl). Pentavalent methylarsenicals are progressively less toxic than inorganic arsenate, while trivalent methylarsenicals are progressively more toxic than inorganic arsenite with the exception of trimethylarsine which is the least toxic arsenic species [4, 5]. Additionally, volatilization of arsenic can occur through methylation [6], which has varied impacts. Methylated forms of arsenic can be released to new areas through air [7], captured during bioremediation [8], or accumulate in crops such as rice [9]. Microbial transformations of arsenic can have consequences for arsenic speciation and methylation; therefore, they impact arsenic ecotoxicity and the fate of arsenic in the environment.

Arsenic biogeochemical cycling by microbial communities is both an ancient [10, 11] and a contemporary [3, 12] phenomenon. Changes to the methylation or oxidation state of arsenic alter biogeochemical cycling of arsenic, and microbes have evolved a variety of mechanisms to carry out these functions. Arsenic related genes are generally separated into two categories: resistance and metabolism [13]. Arsenic resistance, or detoxification, is encoded by the *ars* operon [14]. The *ars* operon protects the cell from arsenic but does not detoxify arsenic itself in the environment. This operon includes arsenite efflux (ArsB, Acr3) which is potentially precluded by cytoplasmic arsenate reduction with either glutaredoxin (ArsC (grx)) or thioredoxin (ArsC (trx)) [14]. Arsenic metabolisms include methylation (ArsM), oxidation (AioAB, ArxAB), and dissimilatory reduction (ArrAB) [13]. While these genetic determinants of arsenic detoxification and metabolism are well-characterized, the full scope of arsenic detoxification and metabolism gene distribution, diversity, and interspecies transfer is unknown [15–17].

Microbial arsenic resistance is reportedly widespread in the environment. Arsenic resistant organisms have been found in sites with low arsenic concentrations (< 7 ppm) [18, 19], and it has been speculated that nearly all organisms have arsenic resistance genes [20]. While the number of identified microorganisms with arsenic resistance genes continues to grow [13], the number of microorganisms without arsenic resistance genes is unclear. Furthermore, though the complete arsenic biogeochemical cycle has been detected in the environment [10], the relative contributions of genes encoding detoxification and metabolism remain unknown [11]. A global, biogeographic perspective of environmental arsenic related genes would improve understanding of their ecology. This information would expand foundational knowledge of arsenic detoxification and metabolism, including local and global abundances, gene diversity, dispersal across different environments, and representations over the microbial tree of life.

Knowledge gaps concerning the diversity of microbial arsenic related genes are driven, in part, by numerous inconsistencies in nomenclature and detection methods. Though public microbial metagenome and genome data continue to surge, there are several practical hurdles to achieving a robust, global assessment of microbial arsenic related genes from this wealth of data. First, tools to detect these genes rely on imperfect annotation [15] and widely vary in nomenclature [21]. Next, the use of different reference databases [12, 22–25] and normalization techniques [25, 26] complicates comparisons between studies. To overcome these hurdles, we developed an open-access toolkit to examine arsenic resistance and metabolism genes in microbial sequence datasets. This toolkit allowed us probe genomic and metagenomic datasets simultaneously to investigate arsenic related genes in soil microbiomes. We first asked whether arsenic related genes are universal in soil-associated microorganisms. Next we tested the hypothesis that genes encoding arsenic detoxification are more abundant than those encoding arsenic metabolism. We also tested the hypothesis that arsenic resistance genes with redundant function (i.e. *acr3* and *arsB; arsC* (grx) and *arsC* (trx)) would have complementary environmental abundances. Third, we asked whether estimations of arsenic related gene abundance are biased by cultivation efforts, as cultivation is often a research emphasis because cultivable, arsenic resistant microorganisms can be used in bioremediation [17]. Finally, we tested the hypothesis that sequence variants of arsenic related genes are endemic, not cosmopolitan.

## Results

### A bioinformatic toolkit for detecting and quantifying arsenic related genes

We developed a toolkit to improve investigations of microbial arsenic related genes (**Figure 1AB**)[14, 27–31]. We selected these nine genes because they are markers of arsenic detoxification and metabolism [21, 25] and because their genetic underpinnings are well established. Seed sequences (high quality and full length sequences) for each gene of interest were collected and used to construct BLAST databases [32], functional gene (FunGene) databases [33], hidden Markov models (HMMs [34]), and gene resources for gene-targeted assembly (Xander [35]) (**Figure 1A**). Altogether, this toolkit relies on consistent references and nomenclature and can search both amino acid and nucleotide sequence data.

**Figure 1.**
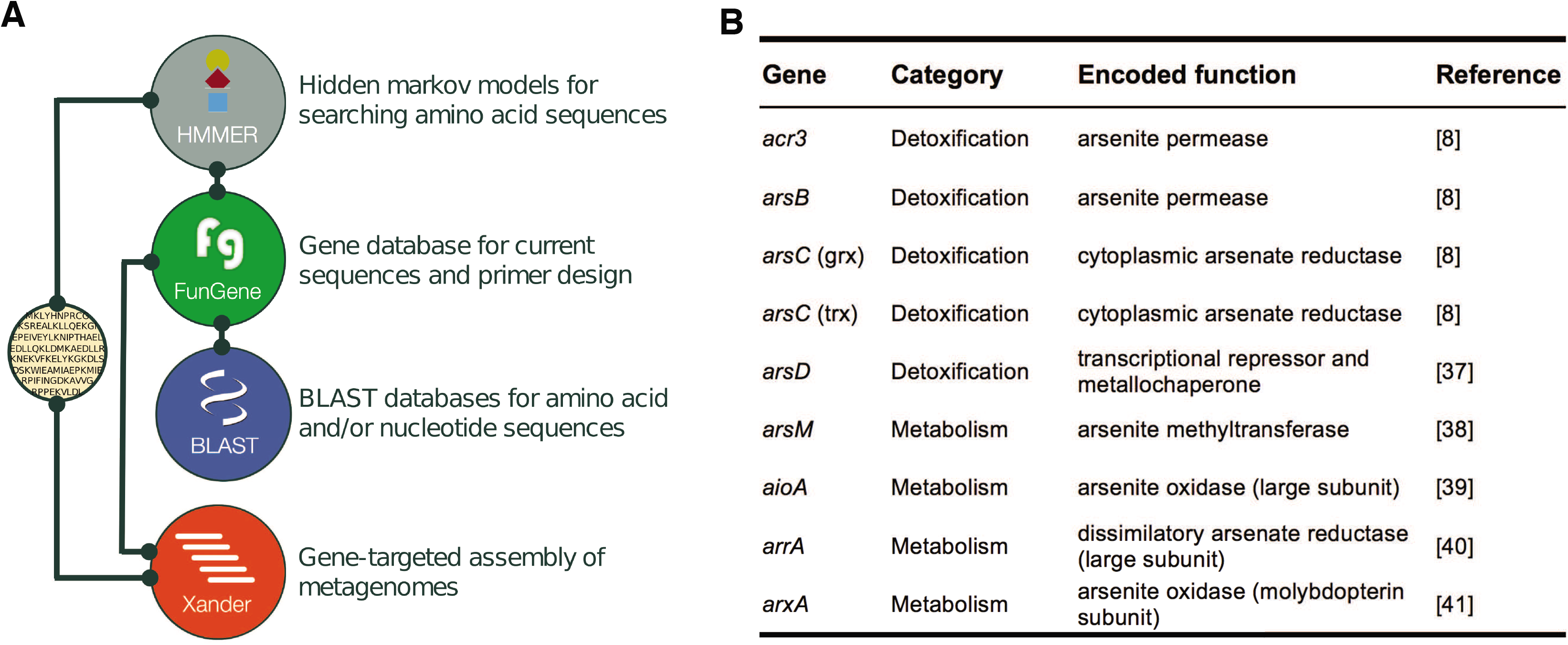
Arsenic resistance and metabolism gene toolkit schematic. **A)** Seed sequences for nine arsenic resistance genes were used to construct an arsenic resistance gene database with existing tools [33, 35, 44, 74]. Lines indicate interdependence between modules. **B)** Table of arsenic resistance and metabolism genes included in the toolkit. The toolkit is freely available on GitHub: github.com/ShadeLab/meta_arsenic

To demonstrate the utility of our toolkit, we performed an analysis of arsenic related genes in soil-associated genomes and metagenomes. We used HMMs for marker genes for arsenic detoxification and metabolism to search RefSoil+ genomes, a set of complete chromosomes and plasmids from cultivable soil microorganisms [36]. Additionally, we used a gene-targeted assembler [35] to test 38 public soil metagenomes from Brazil, Canada, Malaysia, Russia, and the United States for arsenic resistance and metabolism genes (**Additional File 1**). Ultimately, these data serve as a broad baseline of arsenic detoxification and metabolism genes in soil.

### Phylogenetic distributions and genomic locations of arsenic related genes

We asked whether arsenic resistance and metabolism genes were universal in RefSoil+ organisms [36]. Of the 922 RefSoil+ genomes spanning 25 phyla (**Figure 2B; Additional File 2**), 14.3% (132 genomes) did not contain any tested arsenic related genes. Of the 25 phyla in RefSoil+, two phyla (Chlamydiae and Crenarchaeota) did not have any of these genes. These phyla, however, had few RefSoil+ representatives (three and nine, respectively), so other members of these phyla may have arsenic detoxification and metabolism genes. Supporting this hypothesis, a Crenarchaeota isolate was previously reported to oxidize arsenic [37]. Nonetheless, these data suggest that arsenic related genes are widespread, but not universal, even among cultivable soil organisms (**Figure 2**).

**Figure 2.**
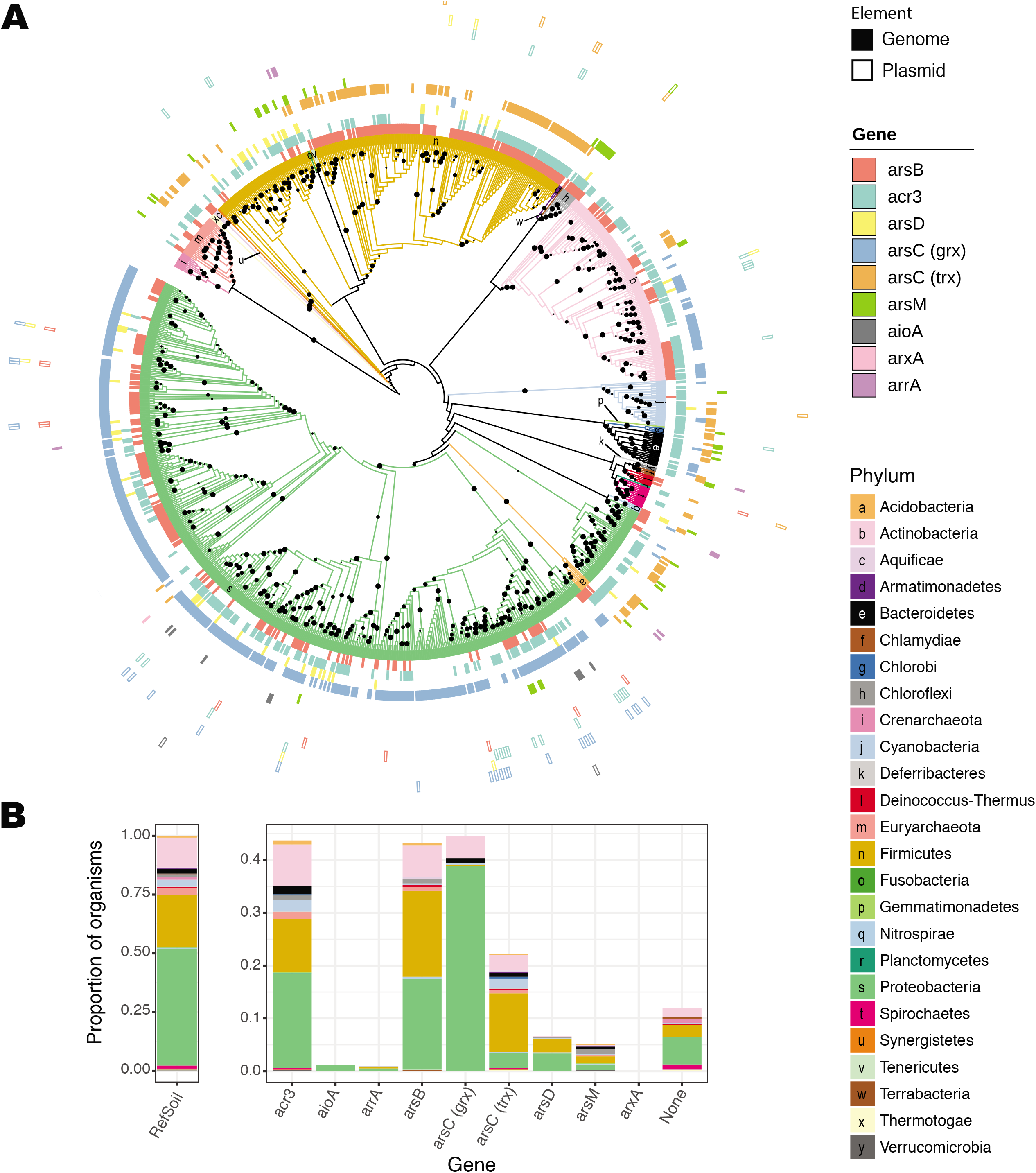
Arsenic resistance and metabolism genes in RefSoil+ organisms. **A)** Maximum likelihood tree of 16S rRNA genes in RefSoil+ organisms. Bootstrap support > 50 is shown with black circles. Tree branches and the first ring are colored by organism taxonomy. Each node is annotated with arsenic resistance genotype where color indicates the gene. Filled boxes indicate gene presence on chromosome, and open boxes indicate gene presence on plasmid. **B)** Proportion of RefSoil+ organisms and organisms containing arsenic resistance genes are colored by the taxonomy of the organism containing the gene. “None” refers to the number of genomes that do not test positive for any of the nine arsenic resistance genes analyzed. Note the difference between y-axes.

We next asked whether 16S rRNA gene phylogeny was predictive of arsenic genotypes using a test for phylogenetic signal (Bloomberg’s K [38]). No phylogenetic signal was observed for plasmid-borne sequences or genes encoding arsenic metabolisms (*aioA, arrA, arxA*); however, relatively few RefSoil+ microorganisms tested positive for these genes. Despite their phylogenetic breadth (**Additional Files 3 – 7**), chromosomally-encoded *acr3, arsB, arsC* (grx), *arsC* (trx), and *arsM* were similar between phylogenetically related organisms (false discovery rate adjusted p < 0.01; **Figure 2A**).

### Phylogenetic diversity of arsenic related genes: insights into vertical and horizontal transfer

#### Arsenite efflux pumps

We examined the phylogenetic diversity of distinct genes encoding arsenite efflux pumps, *acr3* and *arsB*, for soil-associated microorganisms (**Figure 3, Additional Files 3 – 4**). Gene *acr3* is separated into two clades: *acr3*(1) and *acr3*(2) [39]. Clade *acr3*(1) is typically composed of Proteobacterial sequences while *acr3*(2) is typically composed of Firmicutes and Actinobacterial sequences [21, 39, 40]. Though RefSoil+ genomes were mostly composed of *acr3*(2) sequences from Proteobacteria (**Figure 3A; Additional File 3**), we observed greater taxonomic diversity observed than previously reported for this clade [21, 39, 40]. Surprisingly, there were deep branches in *acr3*(2) that belonged to Bacteroidetes, Euryarchaeota, Firmicutes, Fusobacteria, and Verrucomicrobia. Similarly, *acr3*(1) contained closely related *acr3* sequences present in a diverse array of phyla (10 out of 25). Both clades had sequences present on plasmids (6.1%). Plasmid-borne *arsB* sequences were only present in Proteobacteria and Deinococcus-Thermus strains (**Figure 3B; Additional File 4**). Sequences from Actinobacteria, Proteobacteria, and Firmicutes were each present in two distinct phylogenetic groups, and previous studies also observed separation of *arsB* sequences based on phylum [39, 40]. Interestingly, our genome-centric analysis revealed that microorganisms with multiple copies of *arsB* did not harbor identical copies. For example, seven *Bacillus subtilis* subsp. *subtilis* strains had two copies of *arsB*, with one from each of the two clades (**Additional File 4**).

**Figure 3.**
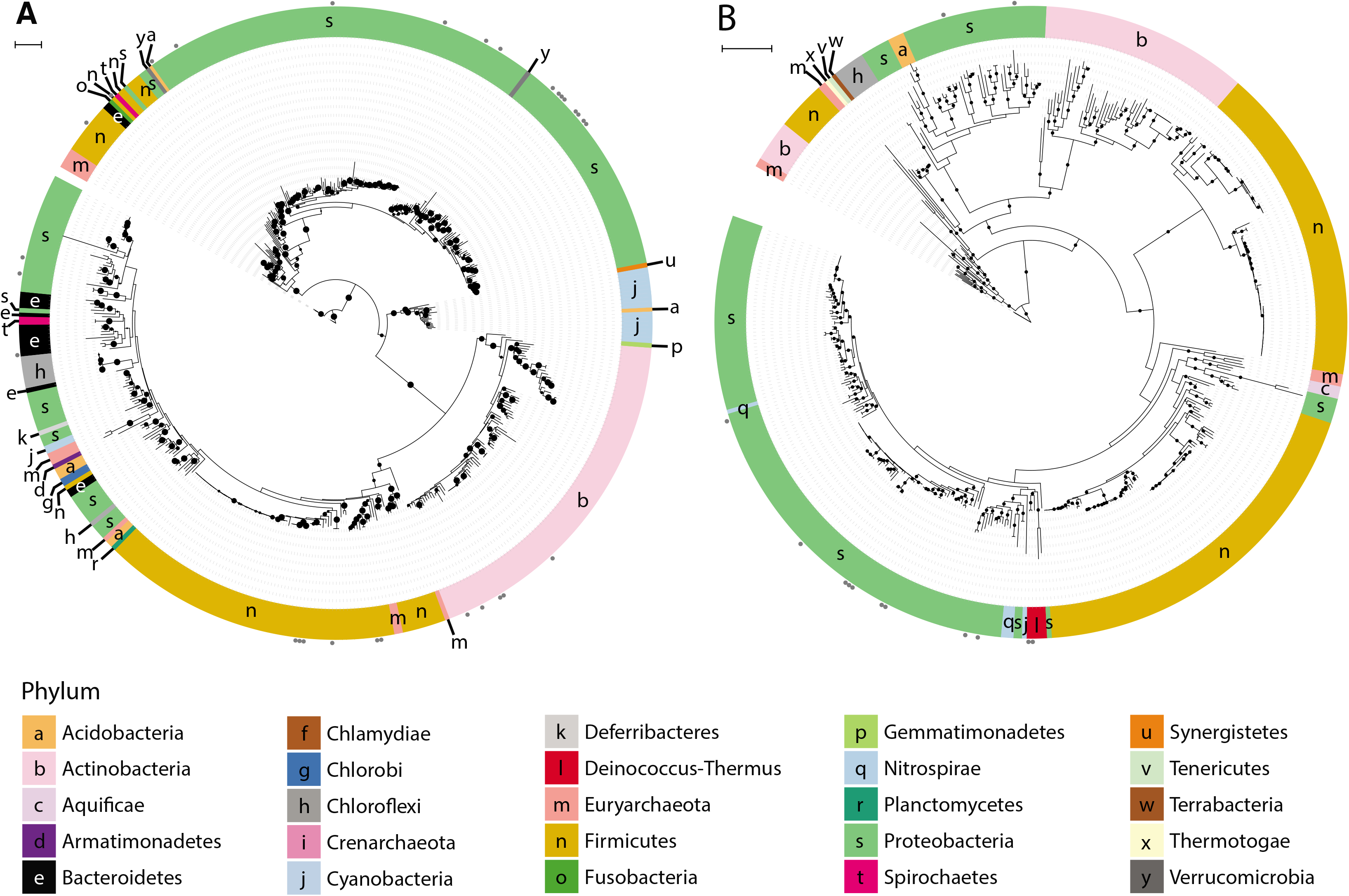
Phylogeny of arsenite efflux pumps in RefSoil+ organisms. Maximum likelihood tree with 100 bootstrap replications of **A)** Acr3 and **B)** ArsB sequences predicted from RefSoil+ genomes. Tree scale = 1. Leaf tip color indicates phylum-level taxonomy. Bootstrap values > 50 are represented by black circles within the tree. Grey circles on the exterior of the tree indicate that a hit was detected on a plasmid and not a chromosome.

#### Cytoplasmic arsenate reductases

Cytoplasmic arsenate reductase (ArsC (trx)) was phylogenetically widespread in RefSoil+ microorganisms (**Figure 4A; Additional File 5**). While some *arsC* (trx) sequences were plasmid-borne, the majority were chromosomally encoded. Similarly, plasmid encoded *arsC* (grx) made up 4.6% of RefSoil+ hits (**Figure 4B; Additional File 6**). Notably, several Proteobacteria strains have multiple copies of *arsC* (grx) with distinct sequences. It is possible that this is the result of an early gene duplication event or HGT of a second *arsC* (grx).

**Figure 4.**
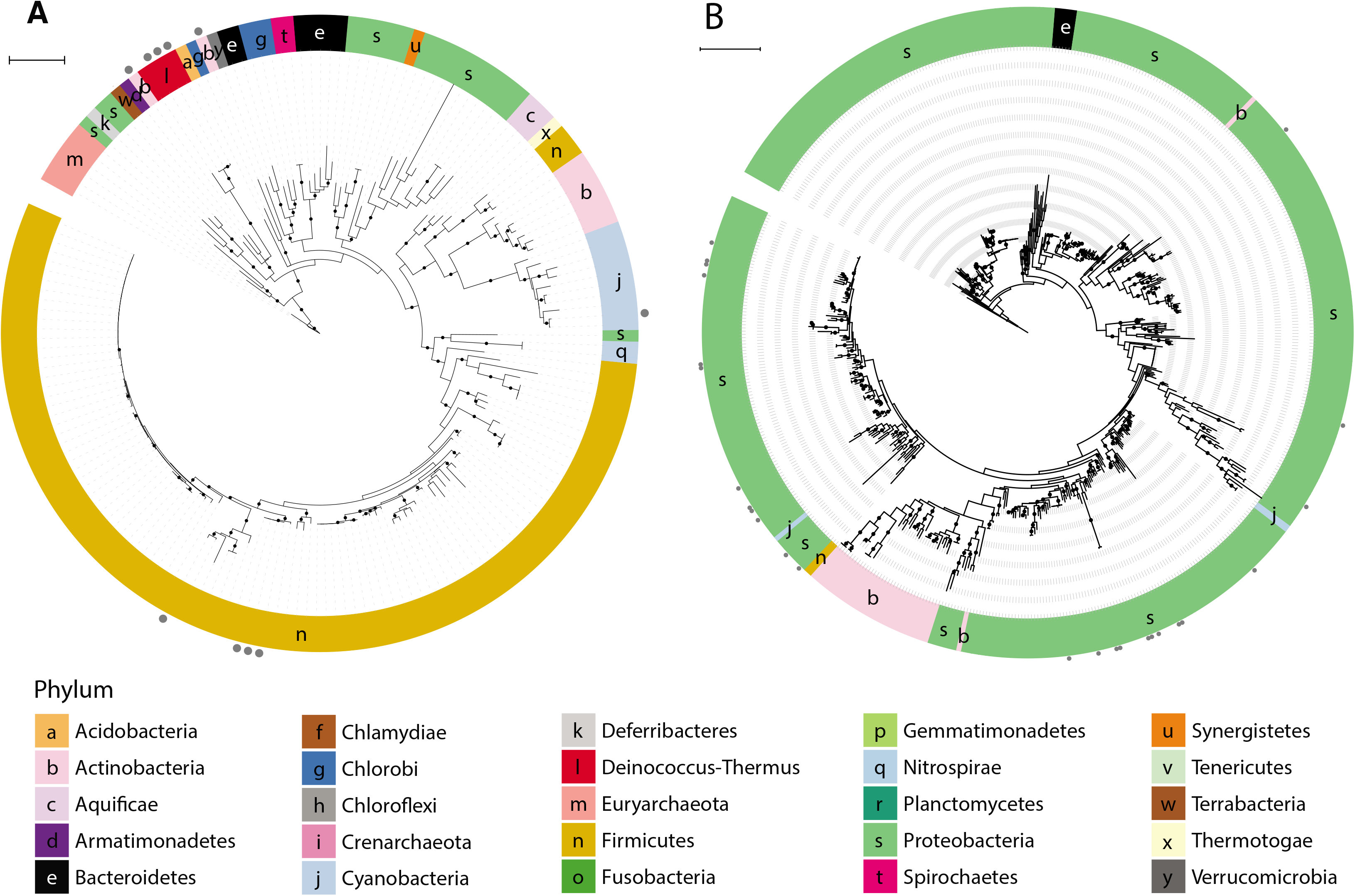
Phylogeny of cytoplasmic arsenate reductases in RefSoil+ organisms. Maximum likelihood tree with 100 bootstrap replications of **A)** ArsC (trx) and **B)** ArsC (grx) sequences predicted from RefSoil+ genomes. Tree scale = 1. Leaf tip color indicates phylum-level taxonomy. Bootstrap values > 50 are represented by black circles within the tree. Grey circles on the exterior of the tree indicate that a hit was detected on a plasmid and not a chromosome.

#### Arsenic metabolisms

*arsM* was relatively uncommon in RefSoil+ microorganisms (5.2%) (**Figure 2**). In the RefSoil+ database, *arsM* was observed in Euryarchaeota as well as several bacterial phyla Acidobacteria, Actinobacteria, Armatimonadetes, Bacteroidetes, Chloroflexi, Cyanobacteria, Firmicutes, Gemmatimonadetes, Nitrospirae, Proteobacteria, Verrucomicrobia (**Figure 5; Additional File 7**). Notably, only one RefSoil+ microorganism, *Rubrobacter radiotolerans* (NZ_CP007516.1), had a plasmid-borne *arsM*.

**Figure 5.**
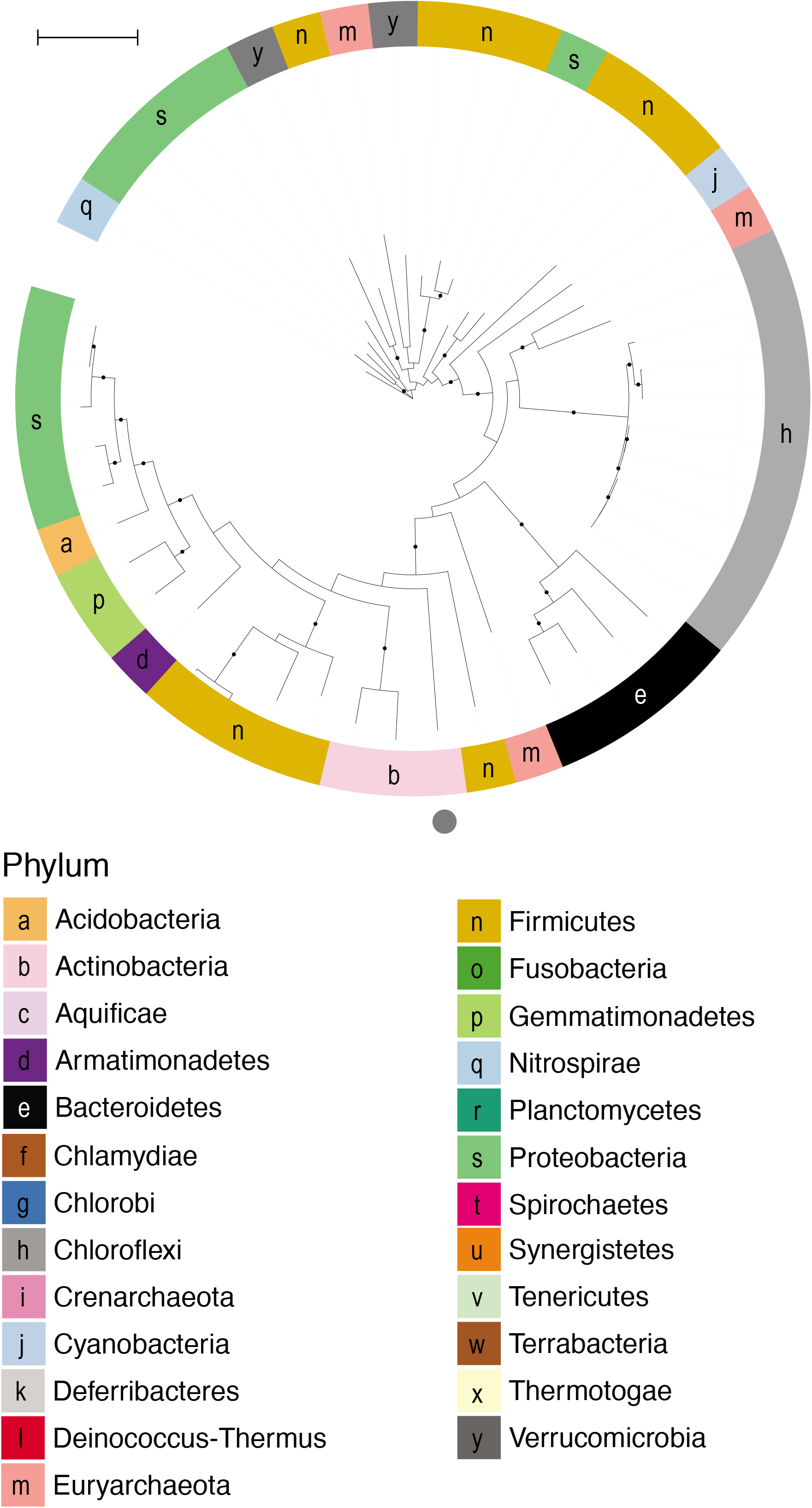
Phylogeny of ArsM in RefSoil+ organisms. Maximum likelihood tree with 100 bootstrap replications of ArsM sequences predicted from RefSoil+ genomes. Tree scale = 1. Leaf tip color indicates phylum-level taxonomy. Bootstrap values > 50 are represented by black circles within the tree. Grey circles on the exterior of the tree indicate that a hit was detected on a plasmid and not a chromosome.

Arsenic metabolism genes *aioA, arrA*, and *arxA* were phylogenetically conserved (**Figure 6**). Genes encoding arsenite oxidases *aioA* and *arxA* were restricted to Proteobacteria. *aioA* sequences clustered into two clades based on class-level taxonomy: all Alphaproteobacteria sequences cluster separately from Gamma- and Betaproteobacteria sequences. The gene encoding dissimilatory arsenate reduction *arrA* was also phylogenetically conserved in RefSoil+ strains, with strains from Proteobacteria clustering separate from Firmicutes (**Figure 6**).

**Figure 6.**
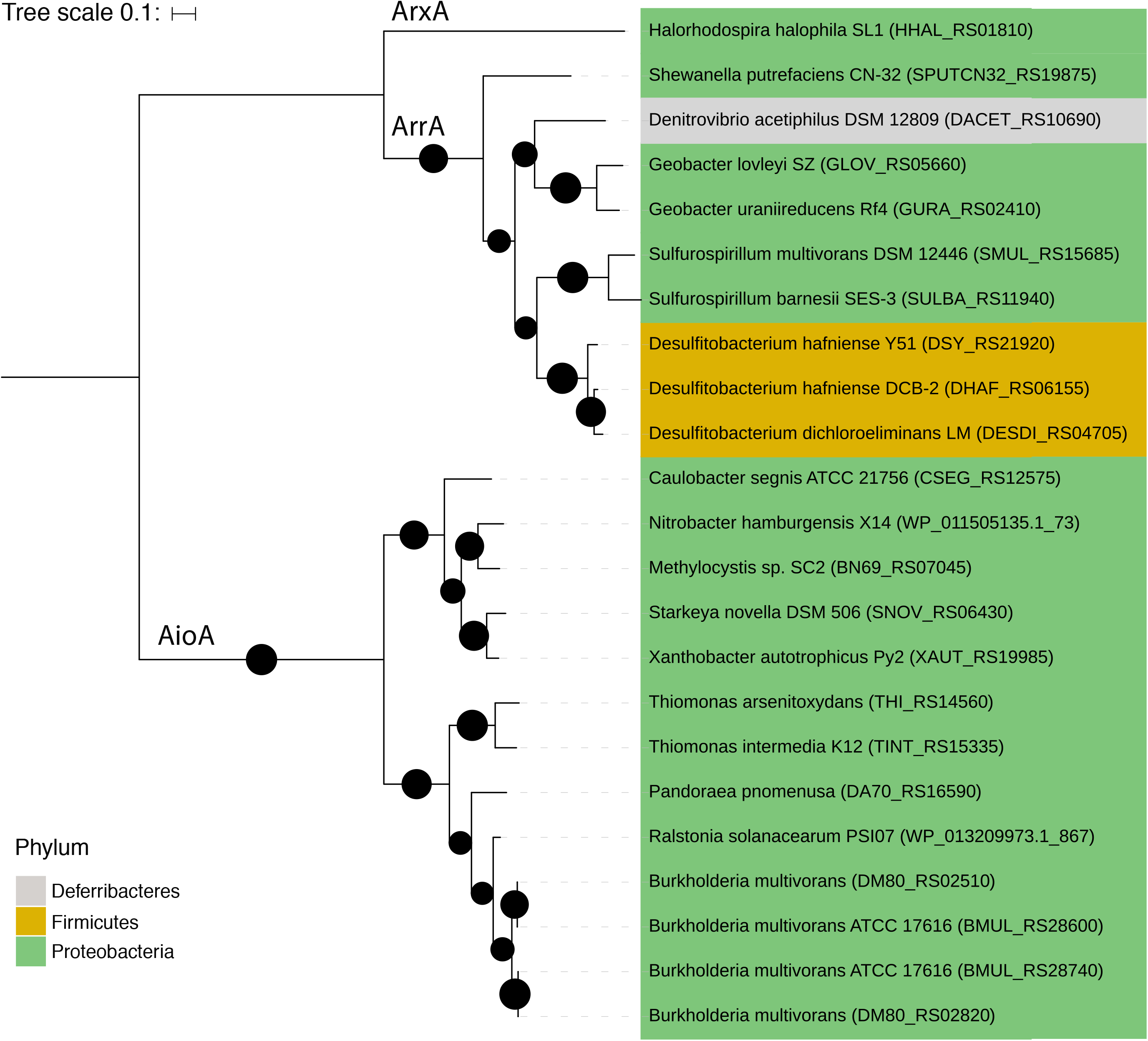
Phylogeny of AioA, ArrA, and ArxA in RefSoil+ organisms. Maximum likelihood tree with 100 bootstrap replications of dissimilatory arsenic resistance proteins predicted from RefSoil+ genomes. Tree scale = 0.1. Leaf tips show the name of the RefSoil+ organisms and background color indicates phylum-level taxonomy. Bootstrap values > 50 are represented by black circles within the tree.

### Cultivation bias and environmental distributions of arsenic related genes

To gain a cultivation-dependent perspective of the abundances of arsenic related genes in soils, we used inferred environmental abundances of RefSoil microorganisms [41, 42]. The environmental abundance of RefSoil microorganisms, which are cultivable, soil-associated microorganisms, was previously estimated by comparing 16S rRNA gene sequences in RefSoil with those in soil metagenomes [41]. We used this estimated abundance of cultivable microorganisms along with arsenic related gene information from this study (**Figure 2**) to estimate the environmental abundances of arsenic related genes from the cultivated bacteria. Arsenic metabolism genes (*aioA, arrA, arsM, arxA*) were predicted to be less common in the environment compared with arsenic detoxification genes (*acr3, arsB, arsC* (grx), *arsC* (trx), and *arsD*) (**Figure 7A**; Mann Whitney U test p < 0.01). Despite similar distributions of *acr3* and *arsB* in RefSoil+ (**Figure 2B**), *acr3* was more abundant in most soil orders (**Figure 7A**; Mann Whitney U test p < 0.05). For genes encoding cytoplasmic arsenate reductases, *arsC* (grx) was more abundant than *arsC* (trx) (Mann Whitney U test p < 0.01).

**Figure 7.**
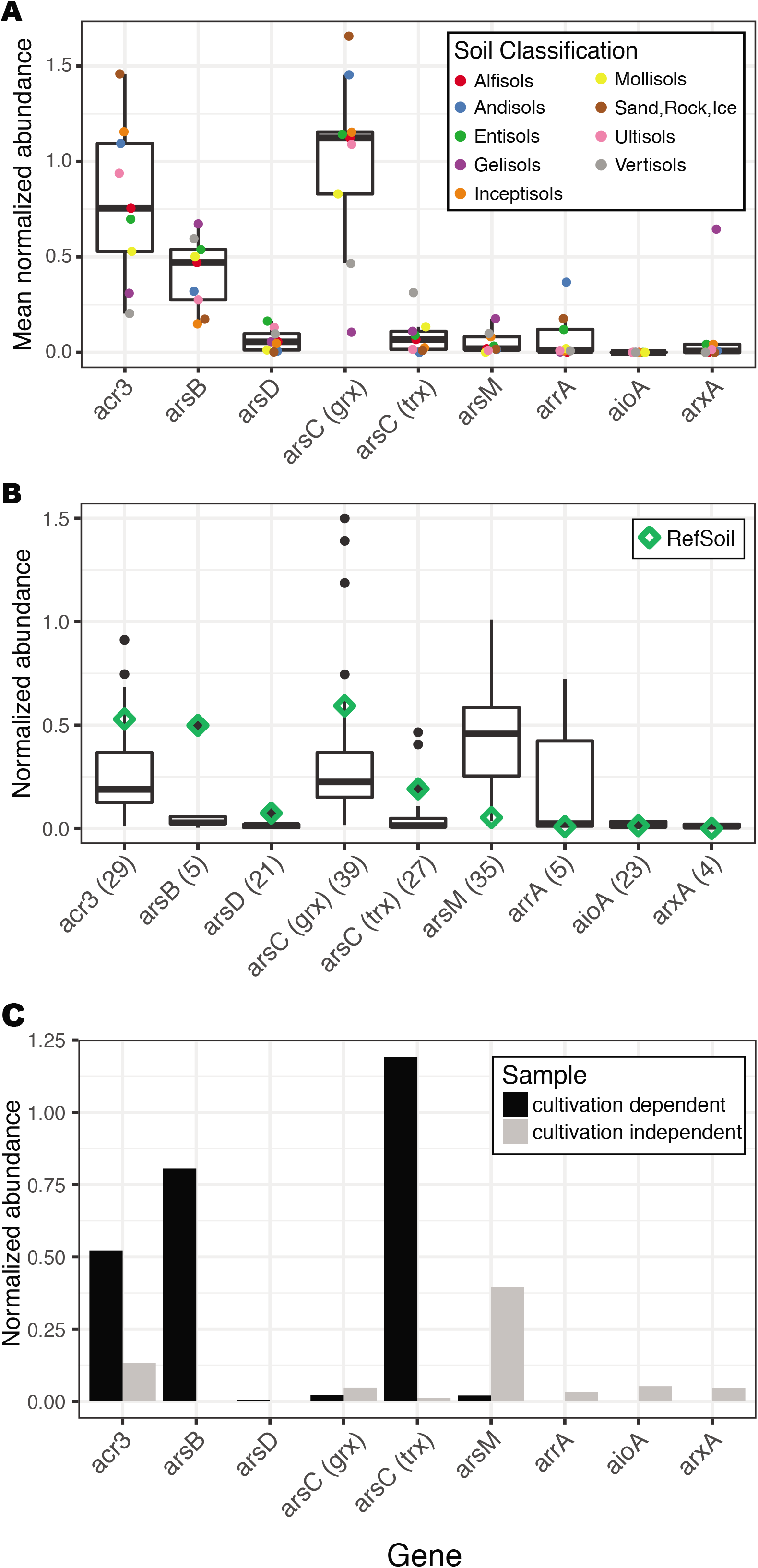
Comparison of arsenic resistance and metabolism gene abundance between cultivation dependent and cultivation independent methods. **A)** Mean normalized abundance of arsenic related genes based on RefSoil microorganisms abundance estimated from corresponding 16S rRNA gene abundance in Earth Microbiome Project datasets. Points are colored by soil order. **B)** Normalized abundance of arsenic resistance genes in RefSoil+ and 38 metagenomes. Metagenome abundance was normalized to *rplB*, and RefSoil+ normalized abundance was calculated using the number of RefSoil+ genomes. Only metagenomes with an arsenic resistance gene detected are shown, and the total number of datasets (including RefSoil+) is shown in parentheses. **C)** *rplB*-normalized abundance of arsenic resistance genes in cultivation dependent and independent metagenomes from the same soil sample.

To gain a cultivation-independent perspective of the abundances of arsenic related genes, we examined their normalized abundance from soil metagenomes (**Figure 7B**). An undetected gene does not confirm absence, so we present a conservative estimate that only includes metagenomes testing positive for a gene. Arsenic detoxification genes (*acr3, arsB, arsC* (grx), *arsC* (trx), and *arsD*) were more abundant than arsenic metabolism genes (*aioA, arrA, arsM*, and *arxA*) (Mann Whitney-U test p < 0.01; **Figure 7B**). Genes encoding arsenite efflux pumps differed in their abundance with *acr3* being more abundant than *arsB* (Mann Whitney U test p < 0.01). We also observed differences in cytoplasmic arsenate reductases: *arsC* (grx) was more abundant than *arsC* (trx) (Mann Whitney U test p < 0.01).

We explored cultivation bias of arsenic related genes with a case study comparing cultivation-dependent (lawn growth on the standard medium TSA50) and - independent communities from the same soil. Genes in the *ars* operon (*acr3, arsB, arsD*, and *arsC* (trx)) were elevated in the cultivation-dependent metagenome (**Figure 7C**). Additionally, arsenic metabolism genes were not detected (*aioA, arrA, arxA*) or in low abundance (*arsM*) in the cultivation-dependent sample; however, all four of these arsenic metabolism genes were detected in the cultivation-independent sample. Though this is a single case study of cultivation-dependent and independent methods, these results recapitulate the general discrepancies between RefSoil+ genomes and soil metagenomes (**Figure 7B**). This bias has important implications for studies focusing on arsenic bioremediation because cultivation-dependent studies could misestimate the potential of microbiomes for arsenic detoxification and metabolism *in situ*.

### Arsenic related gene endemism

Arsenic related genes are globally distributed, but their biogeography is poorly understood. Broadly, arsenic related genes had comparable abundance among different soils (**Figure 7AB**). The relative distributions of distinct arsenic detoxification and metabolism mechanisms in one site, however, are relevant for predicting the impact of microbial communities on the fate of arsenic. To understand site-specific distributions, we explored soil metagenomes from Brazil, Canada, Malaysia, Russia, and the United States (**Additional File 1**). These 16 sites had differences in community membership (**Additional File 9**) and arsenic related gene content (**Figure 8A**). Geographic location was not predictive of arsenic related gene content (Mantel’s r = 0.03493; p > 0.05). Soils had different distributions of arsenic related genes and therefore differed in their potential impact on the biogeochemical cycling of arsenic. While *arsC* (grx) and *arsM* dominated most samples, their relative proportions varied greatly (**Figure 8A**). RefSoil+ data suggests that *arsM* can be found in Verrucomicrobia (100%, n = 2), which is of particular importance for soil metagenomes since Verrucomicrobia are often underestimated with cultivation-dependent methods [43]. The mangrove sample had the most even proportions of arsenic related genes (**Figure 8A**). This distribution was driven by a high abundance of *arsC* (trx) and *arrA*.

**Figure 8.**
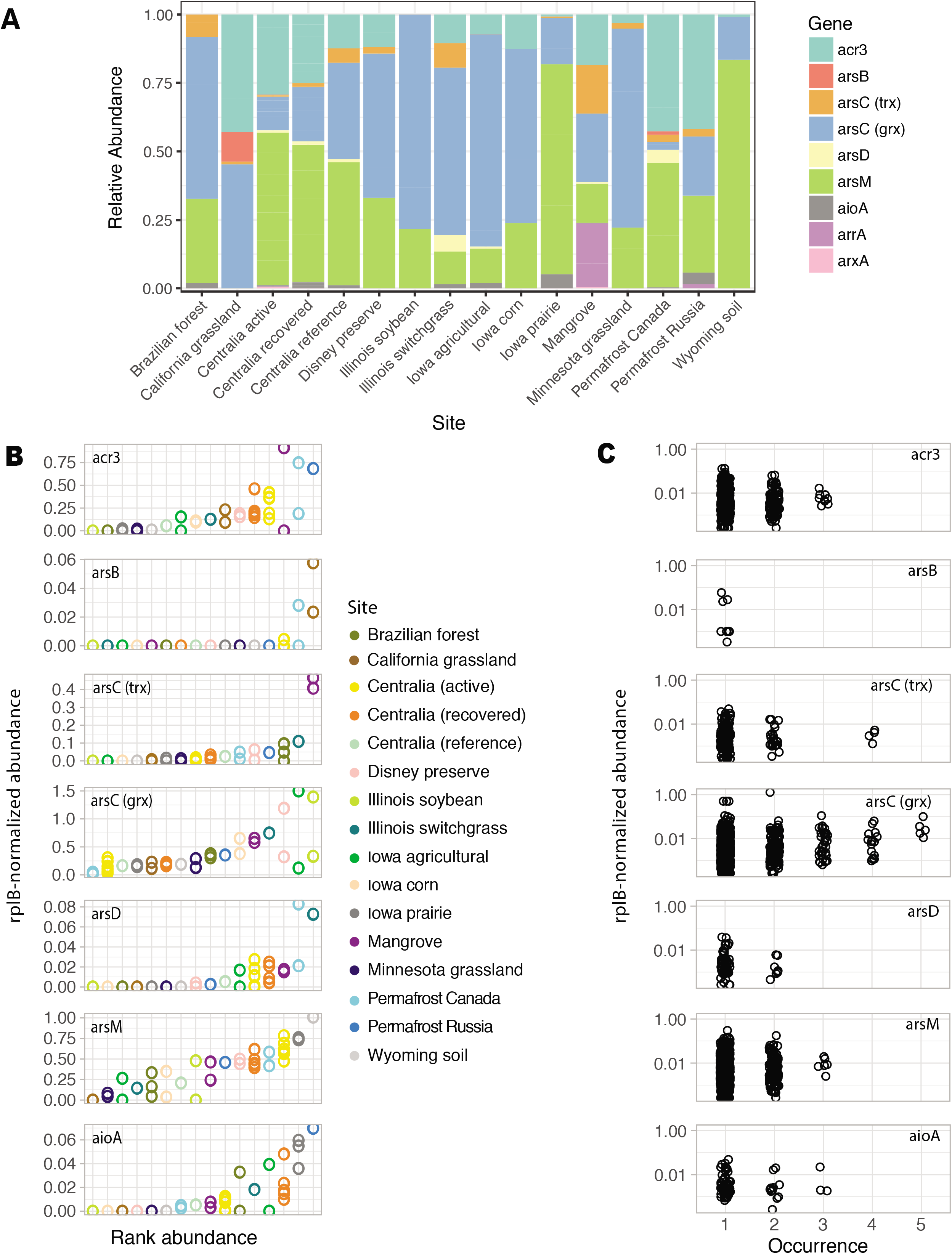
Arsenic resistance and metabolism gene biogeography. **A)** Relative abundance of arsenic resistance genes in soil metagenomes. **B)** Rank *rplB*-normalized abundance of arsenic related genes in soil metagenomes. Sites are ordered by rank mean abundance. Note the differences in y axes. **C)** Abundance-occurrence plots of arsenic related gene sequences clustered at 90% amino acid identity. Number of samples included are as follows Brazilian forest n = 3, California grassland n = 2, Centralia active n = 7, Centralia recovered n = 5, Centralia reference n = 1, Disney preserve n = 2, Illinois soybean n = 2, Illinois switchgrass n = 1, Iowa agricultural n = 2, Iowa corn n = 2, Iowa prairie n = 3, Mangrove n = 2, Minnesota grassland n = 2, Permafrost Canada n = 2, Permafrost Russia n = 1, and Wyoming soil n = 1.

We further examined the arsenic resistance gene abundance at individual sites. We did not include *arr* and *arx* in this analysis due to limited available data. For each gene, the abundance varied greatly, but replicates within one site had similar abundances (**Figure 8B**). The majority of arsenic related gene sequences (99.3%) were endemic and only found in one to two sites, but 24 sequences were detected in three or more sites (**Figure 8C; Additional File 10**). The majority (70.8%) of cosmopolitan sequences belonged to *arsC* (grx). This analysis suggests that arsenic related genes *acr3, arsB, arsC* (trx), *arsD, arsM*, and *aioA* are generally endemic.

## Discussion

### A bioinformatic toolkit for detecting and quantifying arsenic related genes

We developed a toolkit for detecting arsenic related genes from sequence data that supports a variety of applications (**Figure 1A**): arsenic related genes can be detected in amino acid sequences from completed genomes (HMMs [44], BLAST [32]), nucleotide sequences in draft genomes (BLAST), as well as metagenomes and metatranscriptomes (Xander [35]). Because each tool relies on the same seed sequences, there is consistency and opportunity for comparison between sequence datasets that were generated from different sources. While primers already exist for arsenic related genes: *aioA* [45, 46], *acr3* [47], *arsB* [47], *arsC* (grx) [48], *arsC* (trx) [49], *arsM* [9], and *arrA* [50–52], these FunGene [33] databases can be used for testing primer breadth, designing new primers, and browsing sequences.

The toolkit is scalable for additional mechanisms for arsenic resistance and other functional genes of interest (e.g., methylarsenite oxidase (ArsH), C-As lyase (ArsI), trivalent organoarsenical efflux permease (ArsP), organoarsenical efflux permease (ArsJ) [20]), or redox transformations of elements involved in arsenic biogeochemical cycling (e.g., nitrate reductase (NarG) and sulfate reductase (DsrAB) [3, 20]). This toolkit serves as both a resource and an example workflow for developing similar toolkits to examine functional genes, beyond arsenic related genes, in microbial sequence datasets.

### Phylogenetic diversity and distribution of arsenic related genes

It has been conjectured that nearly all organisms have arsenic resistance genes [20], and though this assumption has propagated in the literature, it had never been explicitly quantified. Our data suggest that arsenic detoxification and metabolism genes are ubiquitous, but not universal in RefSoil+ microorganisms (**Figure 2**). It is possible for these 132 organisms to have untested or novel arsenic related genes; nonetheless, these nine well-characterized genes were not universally detected. Additionally, phylogeny was predictive of the presence of *acr3, arsB, arsC* (grc), *arsC* (trx), and *arsM*. This correlation suggests that taxonomy is predictive of arsenic genotype despite documented potential for HGT [19, 39, 53–55]. This result could be explained by ancient rather than contemporary HGT, as seen with *arsM* [55] and *arsC* (grx) [53]. Therefore, we next assessed evidence for HGT by examining the phylogenetic congruence and genomic location (e.g. chromosome or plasmid) of arsenic related gene sequences.

Horizontal transfer of arsenic related genes has been well documented [19, 39, 53–57] and is an important consideration for understanding the propagation and taxonomic identity of arsenic related genes. We examined the phylogenetic diversity of arsenic related genes in RefSoil+ microorganisms, including plasmids and chromosomes, and compared them with the 16S rRNA gene taxonomy.

#### Efflux pumps

While known *acr3* sequences separates into two clades [21, 39, 40], plasmid-borne *acr3* sequences were present across clades, suggesting a potential for transfer across unrelated taxa. Therefore, studies assigning taxonomy to *acr3* in the absence of host information should consider the clade precisely and proceed with caution. Despite their functional redundancy as arsenite efflux pumps, *acr3* and *arsB* have very distinctive diversity. As compared with *acr3, arsB* was less diverse and more phylogenetically conserved (**Figure 3B; Additional File 4**). This observation is in agreement with previous reports comparing the diversity of *arsB* to *acr3* [39, 40]. Multiple, phylogenetically distinct copies of *arsB* were present in some RefSoil+ organisms, which could be due to an early gene duplication and subsequent diversification or to an early transfer event. Therefore, despite relatively lower sequence variation, this *arsB* phylogeny suggests an interesting evolutionary history that could be investigated further.

#### Cytoplasmic arsenate reductases

*arsC* (trx) was predominantly found on RefSoil+ chromosomes, not plasmids, suggesting vertical transfer of *arsC* (trx) is common. *arsC* (trx) was present in both Bacteria and Archaea, and sequences from the two domains formed two distinct clades. *arsC* (trx) sequences that cluster separately from Bacterial-*arsC* (trx) sequences have been documented in Thermococci, Archaeoglobi, Thermoplasmata, and Halobacteria [58] Together, this distribution supports an early evolutionary origin for *arsC* (trx). Thus, *arsC* (trx) appears to be an evolutionarily old enzyme that is phylogenetically conserved despite its presence on plasmids and potential for HGT. Plasmid-encoded *arsC* (grx) were also observed in RefSoil+ microorganisms, highlighting a contemporary potential for HGT that has been documented in soil [53]. Thus, both genes encoding cytoplasmic arsenate reductases were more common on chromosomes.

#### Arsenic metabolisms

The evolutionary history of the gene encoding arsenite S-adenosylmethionine methyltransferase, *arsM*, was recently investigated [54, 55]. Both studies independently determined that *arsM* evolved billions of years ago and was subject to HGT [54, 55]. In this work, *arsM* sequences from Euryarchaeota were dispersed throughout the *arsM* phylogeny, supporting the potential for inter-kingdom transfer events that were recently suggested [54, 55]. Very few RefSoil+ organisms had arsenic metabolism genes *aioA, arrA*, or *arxA*, which limits phylogenetic analysis. Nonetheless, they were mostly found in Proteobacteria, which is in agreement with previous work [13].

### Cultivation bias and environmental distributions of arsenic related genes

Cultivation-based assessments of arsenic related gene content are important since cultivable strains are often favored for bioremediation [59]. We estimated distributions of arsenic related genes in cultivable microorganisms from soils and found a greater abundance of arsenic detoxification genes *acr3, arsB*, and *arsC* (trx) (**Figure 7A**). A previous study also reported an abundance of *acr3* over *arsB* in cultivable microoganisms from forest soils and attributed this to the greater phylogenetic distribution of *acr3* compared with *arsB* [40]. Additionally, they found that *arsC* (grx) was more abundant than *arsC* (trx) in cultivated microorganisms from these soils. It has been posited in cultivation-independent studies that *arsC* (trx) is more efficient than *arsC* (grx) and that high local arsenic concentrations result in a relatively greater abundance of *arsC* (trx) [21, 60]. Our cultivation-dependent abundances suggest that *acr3* and *arsC* (grx), rather than *arsB* and *arsC* (trx), predominantly comprise the arsenic detoxification pathway in soils.

To assess arsenic related gene content without cultivation bias, we examined arsenic related genes in soil metagenomes. As predicted by cultivable organisms, arsenic metabolism genes (*aioA, arrA, arxA*) were generally in low abundance while *acr3* and *arsC* (grx) were in high abundance. Estimates of genes encoding arsenic detoxification (*acr3, arsB, arsD, arsC* (grx), *arsC* (trx)) were considerably lower in these cultivation-independent samples. This result could be due, in part, to the large number of RefSoil+ microorganisms with multiple copies of these genes (**Additional File 8**). Cultivation-independent genomes (e.g., single-cell amplified genomes and metagenome-assembled genomes) could provide greater context about the environmental distributions of copy numbers of arsenic related genes.

Notably, *arsM* was abundant in soil (median 48%), which greatly exceeds cultivation-dependent estimations, and in a case-study of cultivation dependent and independent techniques, *arsM* was more abundant in the cultivation independent sample (**Figure 7C**). Due to the early phylogenetic origins of *arsM* and its independent functionality [55], this abundance of *arsM* in soil metagenomes is not unexpected. *arsM* is typically studied in paddy soils [6, 61, 62], but metagenomes in this study suggest it is an important component of the arsenic biogeochemical cycle in a variety of soils.

### Arsenic related gene endemism

We examined the relative abundance of arsenic related genes in soil metagenomes and observed differences in genetic potential for arsenic transformation that could impact biogeochemical cycling (**Figure 8A**). Notably, the mangrove sample had the most even proportions of arsenic related genes. While the arsenic concentrations in this sample are unknown, mangroves are considered sources and sinks for arsenic [63–65]. This could explain the greater abundance of *arsC* (trx), which is hypothesized to be more abundant in high arsenic sites [21, 60]. Additionally, *arrA* encodes a dissimilatory arsenate reductase that functions in an anaerobic environment [30], so its greater abundance in sediment is expected. Soil geochemical data was not available for all metagenomes examined in this work, so direct comparisons of arsenic related gene content and soil geochemistry were not possible. This highlights the importance and utility of depositing geochemical data with DNA sequences. Future work, however, could further examine relationships between arsenic resistance genes and soil geochemical data, including arsenic concentration and redox potential.

We also measured whether arsenic related gene sequence variants were endemic or cosmopolitan in soil metagenomes (**Figure 8C**). We found that genes *acr3, arsB, arsC* (trx), *arsD, arsM*, and *aioA* were generally endemic, suggesting regional dispersal limitation. Only one *aioA* and three *acr3* sequences were detected in multiple sites. This supports a previous finding that *acr3* and *aioA* from the acid mine drainage in Carnoulès were endemic [66]. Conversely, *arsC* (grx) was cosmopolitan which could suggest genetic migration via HGT or vertical transfer and a limited gene diversification. Both are plausible since *arsC* (grx) was common in RefSoil+ plasmids and had low phylogenetic diversity (**Figure 4B; Additional File 6**).

## Conclusions

We developed a bioinformatic toolkit for detecting arsenic detoxification and metabolism genes in microbial sequence data and applied it to analyze the genomes and metagenomes from soil microorganisms. This toolkit informs hypotheses about the evolutionary histories of these genes (including potential for vertical and horizontal transfers) and how community ecology *in situ* may influence their prevalence and distribution. This study reports the phylogenetic diversity, genomic locations, and biogeography of arsenic related genes in soils, integrating information from different ‘omics datasets and resources to provide a broad synthesis. The toolkit and the synthesis presented here can catalyze future work to understand the ecology and evolution of microbial arsenic biogeochemistry. Furthermore, the toolkit acts as a framework for similar studies of other functional genes of interest.

## Materials and Methods

### Gene Selection and Functional Gene (FunGene) Database Construction

Marker genes can be used to estimate their potential to influence the arsenic biogeochemical cycle [21, 25], so we selected nine well-characterized genes: *acr3, aioA, arsB, arsC* (grx), *arsC* (trx), *arsD, arsM, arrA*, and *arxA*. FunGene databases [33] were constructed for the following arsenic related genes: *arsB, arsC* (grx), *arsC* (trx), *acr3, aioA, arrA*, and *arxA*. The *arxA* database was constructed with seed sequences from [12]. For all other genes, UniProt [67] was used to obtain full length, reviewed sequences when possible. NCBI clusters of orthologous groups (COG) [68] for each gene were examined for evidence of function in the literature. All COG and UniProt sequences were aligned using MUSCLE [69]. Aligned sequences were included in a maximum likelihood tree with 50 bootstrap replications made with MEGA (v7.0,[70]). Sequences that did not cluster with known sequences and had no evidence of function were removed. A final FASTA file for each gene was submitted to the Ribosomal Database Project (RDP) to construct a FunGene database [33]. All arsenic related gene databases are freely available on FunGene (http://fungene.cme.msu.edu/).

### Arsenic related genes in cultivable soil microorganisms

The RefSoil+ database [36] was used to obtain high-quality genomes (chromosomes and plasmids) from soil microorganisms in the Genomes OnLine (GOLD) database [71]. RefSoil+ chromosomes and plasmids were searched with hmmsearch [44] using HMMs from FunGene with an e-value cutoff of 10^−10^. The top hits were analyzed in R [72]. For each gene, scores and percent alignments were plotted to determine quality cutoffs. Stringent percent alignment scores were included since this search was against completed genome sequences: only hits with scores >100 and percent alignment > 90% were included. Hits with the lowest scores were manually examined to test for false positives. Due to false positives, hits against *aioA, arrA*, and *arxA* were further quality filtered to have scores > 1,000. When one open reading frame (ORF) contained multiple hits, the hit with a lower score was removed. Taxonomy was assigned using the RefSoil database [41], and the relative abundance of arsenic related genes within phyla were examined. A 16S rRNA gene maximum likelihood tree of RefSoil+ bacterial strains was constructed with RAxML (v.8.0.6 [73]) based on the Whelan and Goldman (WAG) model with 100 bootstrap replicates (“-m PROTGAMMAWAG -p 12345 -f a -k -x 12345 -# 100”). Based on accession numbers, gene hits were extracted from RefSoil+ sequences and used to construct maximum likelihood trees for each gene.

### Reference Database Construction

Reference gene databases of diverse, near full length sequences were constructed using limited sequences from FunGene databases [33] for the following genes: *acr3, aioA, arrA, arsB, arsC* (grx), *arsC* (trx), *arsD, arsM*, and *arxA*. Seed sequences and hidden Markov models (HMMs) for each gene were downloaded from FunGene, and diverse protein and corresponding nucleotide sequences were selected with gene-specific search parameters (**Additional File 11**). Briefly, minimum amino acid length was set to 70% of the HMM length; minimum HMM coverage was set to 80% as is recommended by Xander software for targeted gene assembly; and a score cutoff was manually selected based on a drop off point. Sequences were de-replicated before being used in subsequent analysis, and final sequence counts are included in **Additional File 11**. Reference databases were converted to publicly available BLAST databases using BLAST+ [74]. Reference and BLAST databases are publicly available on GitHub (https://github.com/ShadeLab/PAPER_Dunivin_meta_arsenic).

### Sample collection and preparation

A soil surface core (20 cm depth and 5.1 cm diameter) was collected in October 2014 from Centralia, PA (GPS coordinates: 40 48.070, 076 20.574). For cultivation-dependent work, a soil slurry was made by vortexing 5 g soil with 25 mL phosphate-buffered saline (PBS) for 1 min. Remaining soil was stored at −80°C until DNA extractions. The soil slurry was allowed to settle for 2 min. 100 μL of the slurry was then removed and serial diluted using PBS to a 10^−2^ dilution. 100 μL of the solution was added to 50% trypticase soy agar (TSA50) with 200 μg/ml cycloheximide to prevent fungal growth. Plates were incubated at 60°C for 72 h. Lawns of growth were extracted by adding 600 μL trypticase soy broth with 25% glycerol to plates. The plate scrapings were stored at −80°C until DNA extraction.

### DNA extraction and metagenome sequencing

DNA for cultivation-independent analysis was manually extracted from soil using a phenol chloroform extraction [75] and the MoBio DNEasy PowerSoil Kit (MoBio, Solana Beach, CA, USA) according the manufacturer’s instructions. DNA extraction for cultivation-dependent analysis was performed in triplicate from 200 μL of plate scrapings using the E.Z.N.A. Bacterial DNA Kit according to the manufacturer’s instructions. All DNA was quantified using a Qubit dsDNA BR Assay Kit (Life Technologies, NY, USA) and was submitted for NGS library prep and sequencing at the Michigan State University Genomics Core sequencing facility (East Lansing, MI, USA). Libraries were prepared using the Illumina TruSeq Nano DNA Library Preparation Kit. After QC and quantitation, the libraries were pooled and loaded on one lane of an Illumina HiSeq 2500 Rapid Run flow cell (v1). Sequencing was performed in a 2 x 150 bp paired end format using Rapid SBS reagents. Base calling was performed by Illumina Real Time Analysis (RTA) v1.18.61 and output of RTA was demultiplexed and converted to FastQ format with Illumina Bcl2Fastq v1.8.4.

### Public soil metagenome acquisition

In total, 38 soil metagenomes were obtained for this work (**Additional File 1**). Datasets from Centralia, PA, were generated in our research group. All other metagenome data sets were obtained from MG-RAST ((http://metagenomics.anl.gov/). The MG-RAST database was searched on May 15, 2017, with the following criteria: material = soil, sequence type = shotgun, public = true. The resulting list of metagenome data sets was ordered by number of base pairs (bp). Metagenomic data sets with the most bp were only included if they were sequenced using Illumina to standardize sequencing errors, had an available FASTQ file for internal quality control, and contained < 30% low quality as determined by MG-RAST. Within high quality Illumina samples, priority for inclusion was given to projects with multiple samples so that comparisons could be made both within and between soil sites. When a project had multiple samples, data sets with the greatest bp were selected. While we prioritized samples with multiple datasets, several replicate samples were omitted early on due to > 30% of data removed during quality filtering, and samples Illinois soil, Russian permafrost, and Wyoming soil have just one sample. This search ultimately yielded 26 data sets from 12 locations and five countries (**Additional File 2**).

### Soil metagenome processing and gene targeted assembly

Sequences from MG-RAST data sets as well as Centralia sample Cen13 were quality controlled using the FASTX toolkit (fastq_quality_filter, “-Q33 -q 30 -p 50”). Twelve datasets from Centralia, PA, were obtained from the Joint Genome Institute and quality filtered as described previously [76]. Quality filtered sequences were used in all downstream analyses. For each data set, a gene targeted metagenome assembler [35] was used to assemble each gene of interest. For each gene of interest, seed sequences, HMMs, and reference gene databases described above were included. For *rplB*, reference gene database, seed sequences, and HMMs from the Xander package were used. In most instances, default assembly parameters were used except to incorporate differences in protein length (i.e. protein is shorter than default 150 amino acids) or to improve quality (i.e. maximum length is increased to improve specificity) (**Additional File 11**). While the assembler includes chimera removal, additional quality control steps were added. Final assembled sequences (operational taxonomic units, OTUs) were searched against the reference gene database as well as the non-redundant database (nr) from NCBI (August 28, 2017) using BLAST [74]. Genes were re-examined if the top hit had an e-value > 10^−5^ or if top hit descriptors were not the target gene. Genes with low quality results were re-assembled with adjusted parameters.

### Soil metagenome comparison

To compare assembled sequences between samples, gene-based OTU tables were constructed. Aligned sequences from each sample were dereplicated and clustered at 90 amino acid identity using the RDP Classifier [77]. Dereplicated, clustered sequences were converted into OTU tables with coverage-adjusted abundance. These tables were subsequently analyzed in R [72]. RplB OTUs were used to compare community structure. The six most abundant phyla were extracted to include at least 75% of each community; the full community structure is available. To compare the abundance of arsenic related genes among data sets, total counts of *rplB* were used to normalize the abundance of each OTU. Relative abundance of arsenic related genes was also calculated for each sample.

## Declarations

### Ethics approval and consent to participate

Not applicable.

### Consent to publish

Not applicable.

### Availability of data and materials

The full arsenic related gene toolkit (BLAST databases, hidden Markov models, and gene resources for Xander) is publicly available on GitHub (https://github.com/ShadeLab/PAPER_Dunivin_meta_arsenic) [78]. Cultivation dependent and cultivation independent Centralia metagenomes from this study are available on NCBI under BioProject PRJNA492298 [79]. All other metagenomes are publicly available, including those from Brazilian forest [80], California grassland [81], Centralia [82], Disney preserve [83], Illinois soybean [84], Illinois switchgrass [85], Iowa agricultural [86], Iowa corn [87], Iowa prairie [88], Mangrove [89], Minnesota grassland [90], Permafrost Canada [91], Permafrost Russia [92], and Wyoming soil [93]. All RefSoil+ and metagenome analyses from this work are also available on GitHub in the HMM_search and gene_targeted_assembly directories respectively.

### Authors’ contributions

TKD and AS designed the study, TKD and SY contributed code and performed the analysis, TKD and AS wrote the manuscript, all authors read and approved the manuscript.

## Acknowledgements

This work was supported by Michigan State University with computing resources provided by the Michigan State Institute for Cyber-Enabled Research. We thank the Ribosomal Database Project for their advice on reference gene database construction.

## Additional files legends

**Additional File 1**. Available metadata and accession numbers for soil metagenomes used in this study.

**Document Name**: Additional_File_1.docx
**Document Format**: Word
**Document Size**: 23 KB

**Additional File 2**. Phylum-level summary of arsenic related genes in RefSoil+ chromosomes and plasmids.

**Document Name**: Additional_File_2.docx
**Document Format**: Excel
**Document Size**: 15 KB

**Additional File 3. Phylogeny of Acr3 in RefSoil+ organisms**. Maximum likelihood tree with 100 bootstrap replications of Acr3 sequences predicted from RefSoil+ genomes. Leaf tips show the name of the RefSoil+ organisms and background color indicates phylum-level taxonomy. Bootstrap values > 50 are represented by black circles within the tree.

**Document Name**: Additional_File_3.png
**Document Format**: PNG
**Document Size**: 4.5 MB

**Additional File 4. Phylogeny of ArsB in RefSoil+ organisms**. Maximum likelihood tree with 100 bootstrap replications of ArsB sequences predicted from RefSoil+ genomes. Leaf tips show the name of the RefSoil+ organisms and background color indicates phylum-level taxonomy. Bootstrap values > 50 are represented by black circles within the tree.

**Document Name**: Additional_File_4.png
**Document Format**: PNG
**Document Size**: 9.6 MB

**Additional File 5. Phylogeny of ArsC (trx) in RefSoil+ organisms**. Maximum likelihood tree with 100 bootstrap replications of ArsC (trx) sequences predicted from RefSoil+ genomes. Leaf tips show the name of the RefSoil+ organisms and background color indicates phylum-level taxonomy. Bootstrap values > 50 are represented by black circles within the tree.

**Document Name**: Additional_File_5.png
**Document Format**: PNG
**Document Size**: 2 MB

**Additional File 6. Phylogeny of ArsC (grx) in RefSoil+ organisms**. Maximum likelihood tree with 100 bootstrap replications of ArsC (grx) sequences predicted from RefSoil+ genomes. Leaf tips show the name of the RefSoil+ organisms and background color indicates phylum-level taxonomy. Bootstrap values > 50 are represented by black circles within the tree.

**Document Name**: Additional_File_6.png
**Document Format**: PNG
**Document Size**: 4.9 MB

**Additional File 7. Phylogeny of ArsM in RefSoil+ organisms**. Maximum likelihood tree with 100 bootstrap replications of ArsM sequences predicted from RefSoil+ genomes. Leaf tips show the name of the RefSoil+ organisms and background color indicates phylum-level taxonomy. Bootstrap values > 50 are represented by black circles within the tree.

**Document Name**: Additional_File_7.eps
**Document Format**: EPS
**Document Size**: 6.2 MB

**Additional File 8. Histogram of arsenic related gene copy numbers in RefSoil+ organisms**. Total copy number is based on hits from both chromosomes and plasmids from the same organism.

**Document Name**: Additional_File_8.eps
**Document Format**: EPS
**Document Size**: 25 KB

**Additional File 9. Phylum-level community structure of soil metagenomes in this study**.

**Document Name**: Additional_File_9.eps
**Document Format**: EPS
**Document Size**: 67 KB

**Additional File 10**. Summary of endemic arsenic related gene sequences. A sequence was considered endemic if it was present in less than three different soil sites.

**Document Name**: Additional_File_10.docx
**Document Format**: Word
**Document Size**: 45 KB

**Additional File 11**. Summary of reference arsenic resistance and metabolism gene sequences from FunGene databases.

**Document Name**: Additional_File_11.docx
**Document Format**: Word
**Document Size**: 53 KB

