## Supplemental Materials: Additional Files and Figures for "A global survey of arsenic related genes in soil microbiomes"

| Project Name | Sample Location | Country | Sample Shortname | Accession location | Project ID | Sample Name | Gbp |
|---|---|---|---|---|---|---|---|
| **ARMO** | Rondonia | Brazil | Brazilian_forest | MG-RAST | mgp3731 | mgm4546395.3 | 13.27 |
| **ARMO** | Rondonia | Brazil | Brazilian_forest | MG-RAST | mgp3731 | mgm4536139.3 | 9.04 |
| **ARMO** | Rondonia | Brazil | Brazilian_forest | MG-RAST | mgp3731 | mgm4535554.3 | 9.69 |
| **Axel Heiberg Permafrost: Part 4A** | Central Axel Heiberg Island | Canada | Permafrost_Canada | MG-RAST | mgp252 | mgm4523023.3 | 6.52 |
| **Axel Heiberg Permafrost: Part 4A** | Central Axel Heiberg Island | Canada | Permafrost_Canada | MG-RAST | mgp252 | mgm4523145.3 | 5.52 |
| **CedarCreek_minsoil_June2013** | Bethel, MN | USA | Minnesota_grassland | MG-RAST | mgp5588 | mgm4541646.3 | 10.65 |
| **CedarCreek_minsoil_June2013** | Bethel, MN | USA | Minnesota_grassland | MG-RAST | mgp5588 | mgm4541645.3 | 9.77 |
| **Fermi-syntheticlongreads** | Fermi National Accelerator Laboratory | USA | Illinois_switchgrass | MG-RAST | mgp14596 | mgm4653791.3 | 7.95 |
| **GED prairie unassembled** | Iowa | USA | Iowa_prairie | MG-RAST | mgp6377 | mgm4539575.3 | 18.79 |
| **GED prairie unassembled** | Iowa | USA | Iowa_prairie | MG-RAST | mgp6377 | mgm4539572.3 | 17.58 |
| **GED prairie unassembled** | Iowa | USA | Iowa_prairie | MG-RAST | mgp6377 | mgm4539576.3 | 17.43 |
| **GP corn unassembled** | Iowa | USA | Iowa_corn | MG-RAST | mgp6368 | mgm4539522.3 | 8.19 |
| **GP corn unassembled** | Iowa | USA | Iowa_corn | MG-RAST | mgp6368 | mgm4539523.3 | 8.12 |
| **Hofmockel Soil Aggregate COB KBASE** | Boone County, IA | USA | Iowa_agricultural | MG-RAST | mgp2592 | mgm4509400.3 | 24.98 |
| **Hofmockel Soil Aggregate COB KBASE** | Boone County, IA | USA | Iowa_agricultural | MG-RAST | mgp2592 | mgm4509401.3 | 7.86 |
| **ISA-SMC-2011** | Auburn, IL | USA | Illinois_soybean | MG-RAST | mgp2076 | mgm4502542.3 | 12.54 |
| **ISA-SMC-2011** | Auburn, IL | USA | Illinois_soybean | MG-RAST | mgp2076 | mgm4502540.3 | 10.60 |
| **Loma_Ridge_grassland** | Loma Ridge, CA | USA | California_grassland | MG-RAST | mgp1992 | mgm4511115.3 | 6.50 |
| **Loma_Ridge_grassland** | Loma Ridge, CA | USA | California_grassland | MG-RAST | mgp1992 | mgm4511062.3 | 5.77 |
| **Mining of new genes and pathways from soil of mangrove forest** | Matang Mangrove Forest | Malaysia | Mangrove | MG-RAST | mgp11628 | mgm4603402.3 | 24.38 |
| **Mining of new genes and pathways from soil of mangrove forest** | Matang Mangrove Forest | Malaysia | Mangrove | MG-RAST | mgp11628 | mgm4603270.3 | 24.54 |
| **NEON** | Disney Wilderness Preserve, FL | USA | Disney_preserve | MG-RAST | mgp13948 | mgm4664918.3 | 11.20 |
| **NEON** | Disney Wilderness Preserve, FL | USA | Disney_preserve | MG-RAST | mgp13948 | mgm4664925.3 | 4.14 |

| | | | | | | | |
|---|---|---|---|---|---|---|---|
| **Permafrost sediments, North-East Siberia, Kolyma lowland** | Kolyma river lowland | Russia | Permafrost_Russia | MG-RAST | mgp7176 | mgm4546813.3 | 19.20 |
| **Ungulate Exclosure 2015** | Wyoming | USA | Wyoming_soil | MG-RAST | mgp15600 | mgm4670120.3 | 6.41 |
| **Surface soil microbial communities from Centralia Pennsylvania** | Centralia, PA | USA | Centralia_recovered | JGI | Gp0112853 | Cen01 | 23 |
| **Surface soil microbial communities from Centralia Pennsylvania** | Centralia, PA | USA | Centralia_recovered | JGI | Gp0112853 | Cen03 | 26 |
| **Surface soil microbial communities from Centralia Pennsylvania** | Centralia, PA | USA | Centralia_recovered | JGI | Gp0112853 | Cen04 | 25 |
| **Surface soil microbial communities from Centralia Pennsylvania** | Centralia, PA | USA | Centralia_recovered | JGI | Gp0112853 | Cen05 | 25 |
| **Surface soil microbial communities from Centralia Pennsylvania** | Centralia, PA | USA | Centralia_fire-affected | JGI | Gp0112853 | Cen06 | 22 |
| **Surface soil microbial communities from Centralia Pennsylvania** | Centralia, PA | USA | Centralia_recovered | JGI | Gp0112853 | Cen07 | 21 |
| **Surface soil microbial communities from Centralia Pennsylvania** | Centralia, PA | USA | Centralia_fire-affected | JGI | Gp0112853 | Cen10 | 36 |
| **Surface soil microbial communities from Centralia Pennsylvania** | Centralia, PA | USA | Centralia_fire-affected | JGI | Gp0112853 | Cen12 | 24 |
| **Surface soil microbial communities from Centralia Pennsylvania** | Centralia, PA | USA | Centralia_fire-affected | JGI | Gp0112853 | Cen14 | 24 |
| **Surface soil microbial communities from Centralia Pennsylvania** | Centralia, PA | USA | Centralia_fire-affected | JGI | Gp0112853 | Cen15 | 20 |
| **Surface soil microbial communities from Centralia Pennsylvania** | Centralia, PA | USA | Centralia_fire-affected | JGI | Gp0112853 | Cen16 | 51 |
| **Surface soil microbial communities from Centralia Pennsylvania** | Centralia, PA | USA | Centralia_reference | JGI | Gp0112853 | Cen17 | 24 |
| **Surface soil microbial communities from an active vent of coal mine fire in Centralia Pennsylvania, Sep 20 ‘18** | Centralia, PA | USA | Centralia_fire-affected | NCBI | SRR7882662 | Cen13 | 56 |

| PHYLUM | CHROMOSOME | | | | | | | | PLASMID | | | | | |
|---|---|---|---|---|---|---|---|---|---|---|---|---|---|---|
| | arsB | acr3 | arsC (grx) | arsC (trx) | arsM | aioA | arxA | arrA | arsB | acr3 | arsC (grx) | arsC (trx) | arsM | aioA |
| ACIDOBACTERIA (7) | 57.1 | 100 | 0 | 14.3 | 14.3 | 0 | 0 | 0 | 0 | 0 | 0 | 0 | 0 | 0 |
| ACTINOBACTERIA (118) | 44.1 | 61.9 | 33.1 | 5.9 | 1.7 | 0 | 0 | 0 | 0 | 4.2 | 0 | 0.8 | 0.8 | 0 |
| BACTEROIDETES (19) | 0 | 73.7 | 31.6 | 36.8 | 21.1 | 0 | 0 | 0 | 0 | 5.3 | 0 | 0 | 0 | 0 |
| CHLOROFLEXI (9) | 88.9 | 88.9 | 0 | 0 | 100 | 0 | 0 | 0 | 0 | 0 | 0 | 0 | 0 | 0 |
| CYANOBACTERIA (26) | 3.8 | 76.9 | 11.5 | 34.6 | 3.8 | 0 | 0 | 0 | 0 | 0 | 0 | 3.8 | 0 | 0 |
| DEINOCOCCUS-THERMUS (6) | 50 | 0 | 0 | 16.7 | 0 | 0 | 0 | 0 | 16.7 | 0 | 0 | 16.7 | 0 | 0 |
| FIRMICUTES (207) | 65.7 | 44.4 | 1 | 45.9 | 5.8 | 0 | 0 | 1.4 | 0 | 3.4 | 0 | 1.9 | 0 | 0 |
| PROTEOBACTERIA (531) | 33.1 | 40.7 | 79.5 | 4 | 1.9 | 2.4 | 0.2 | 0.9 | 2.1 | 4.1 | 5.6 | 0 | 0 | 0.4 |
| SPIROCHAETES (12) | 0 | 25 | 0 | 16.7 | 0 | 0 | 0 | 0 | 0 | 0 | 0 | 0 | 0 | 0 |

Additional File 3

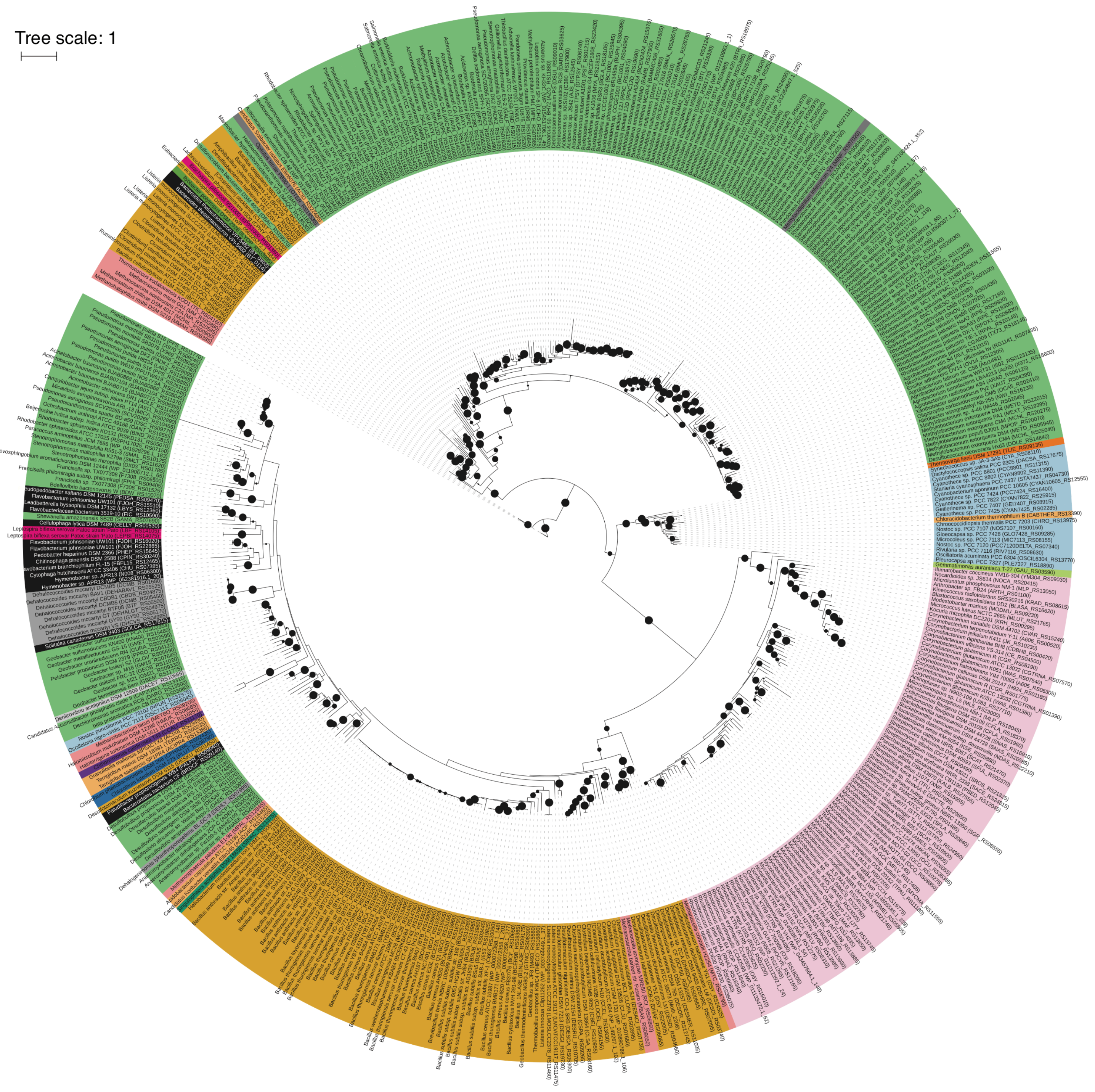

Additional File 4

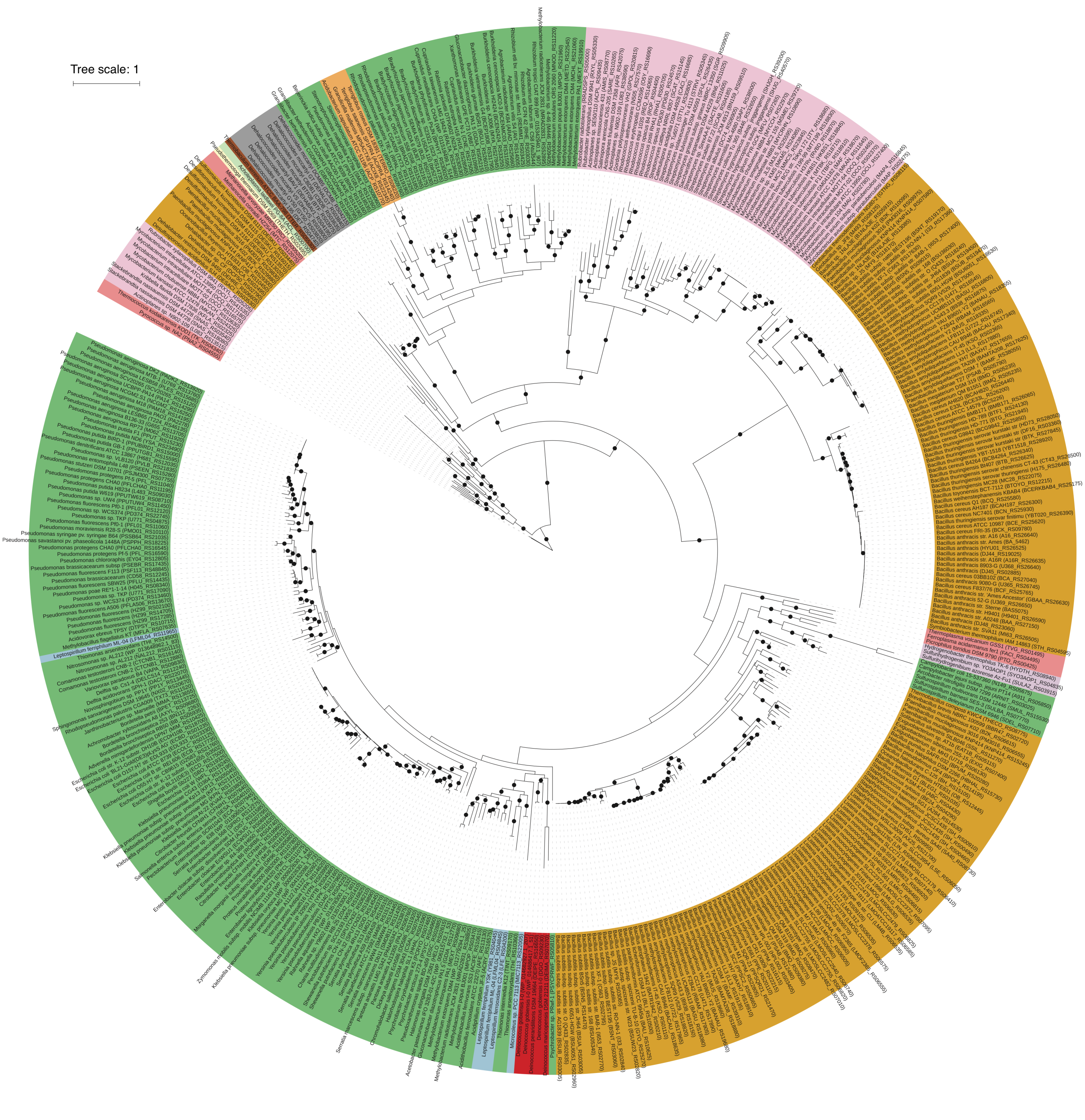

Additional File 5

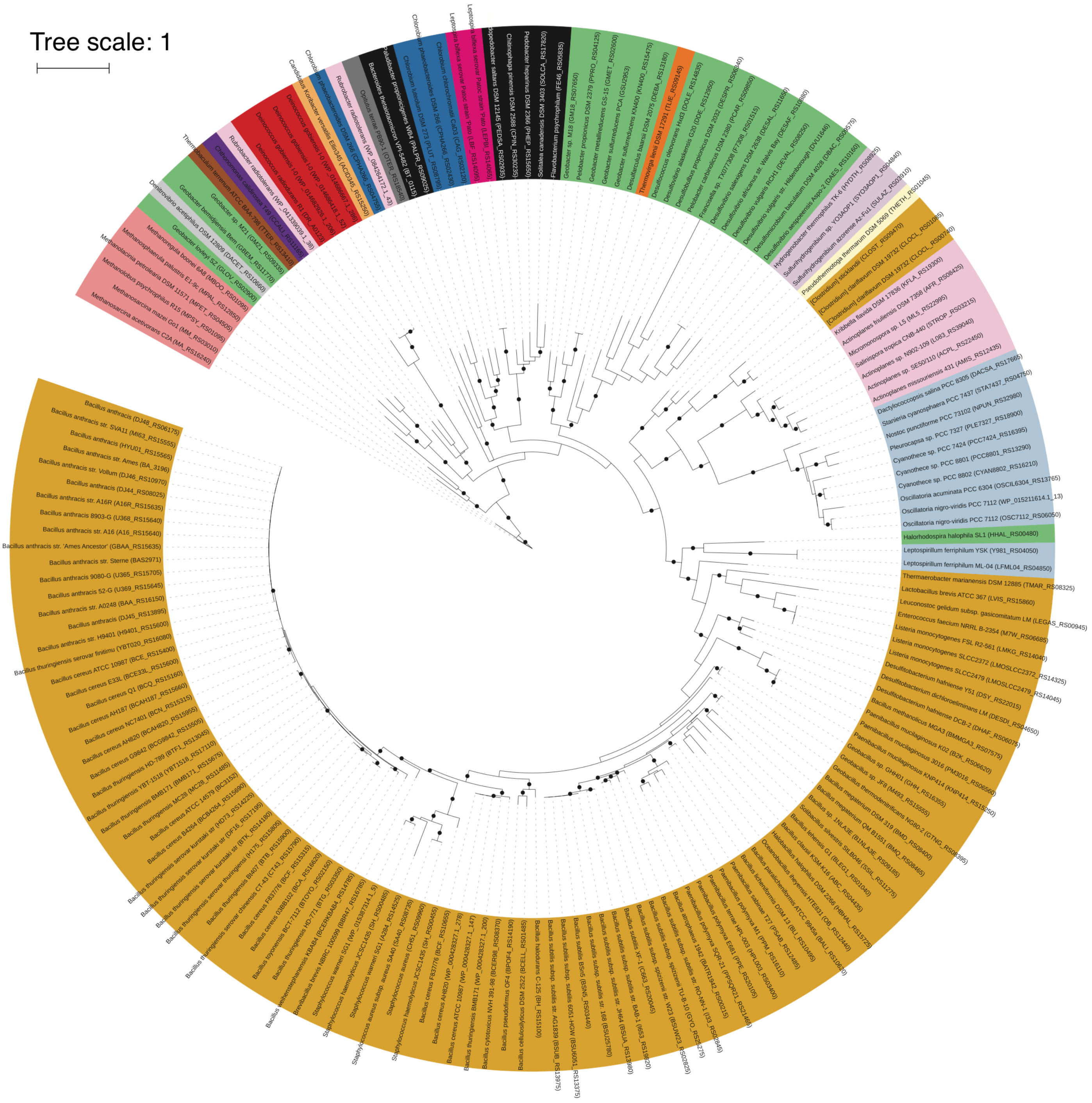

Additional File 6

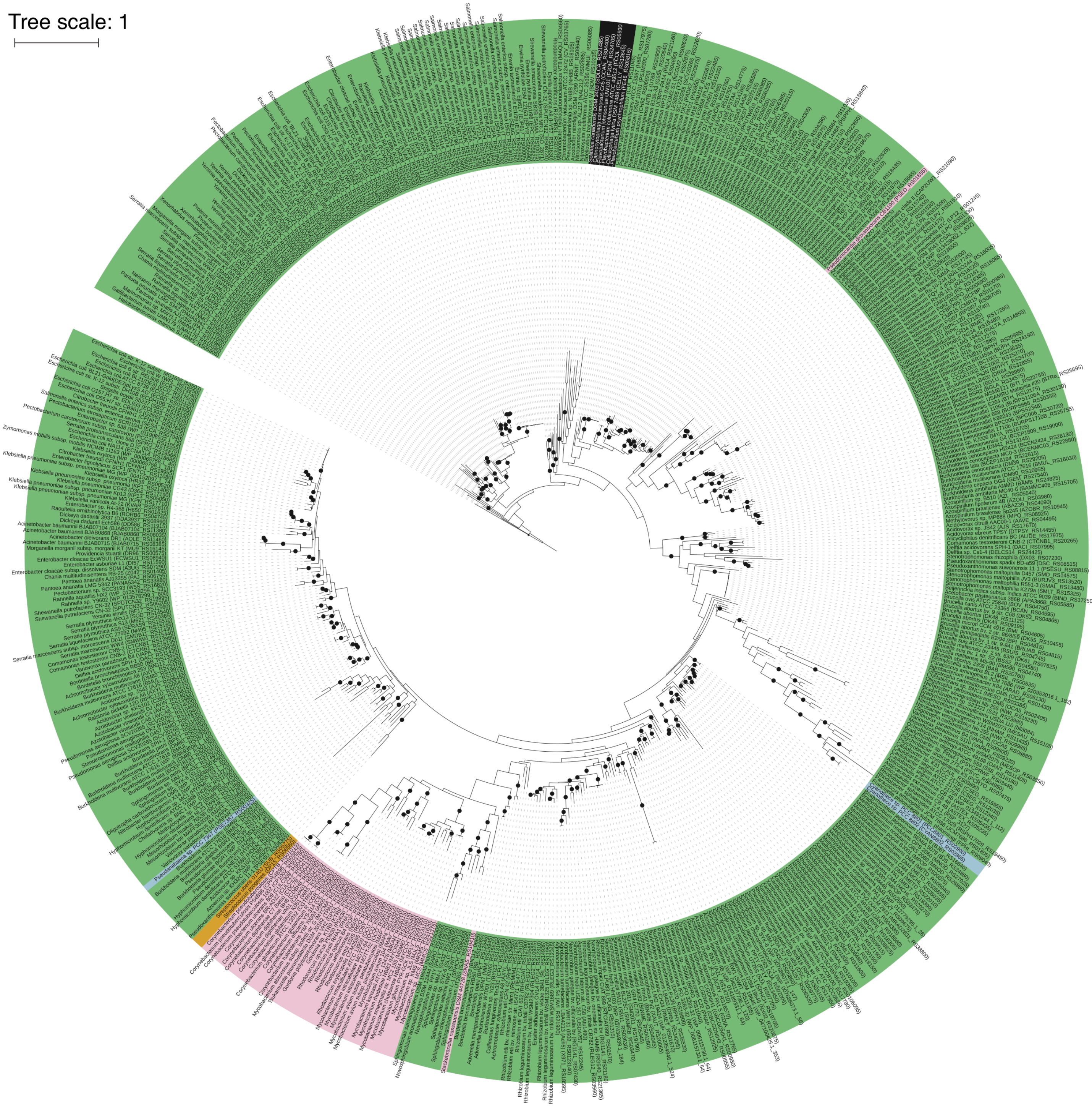

Additional File 7

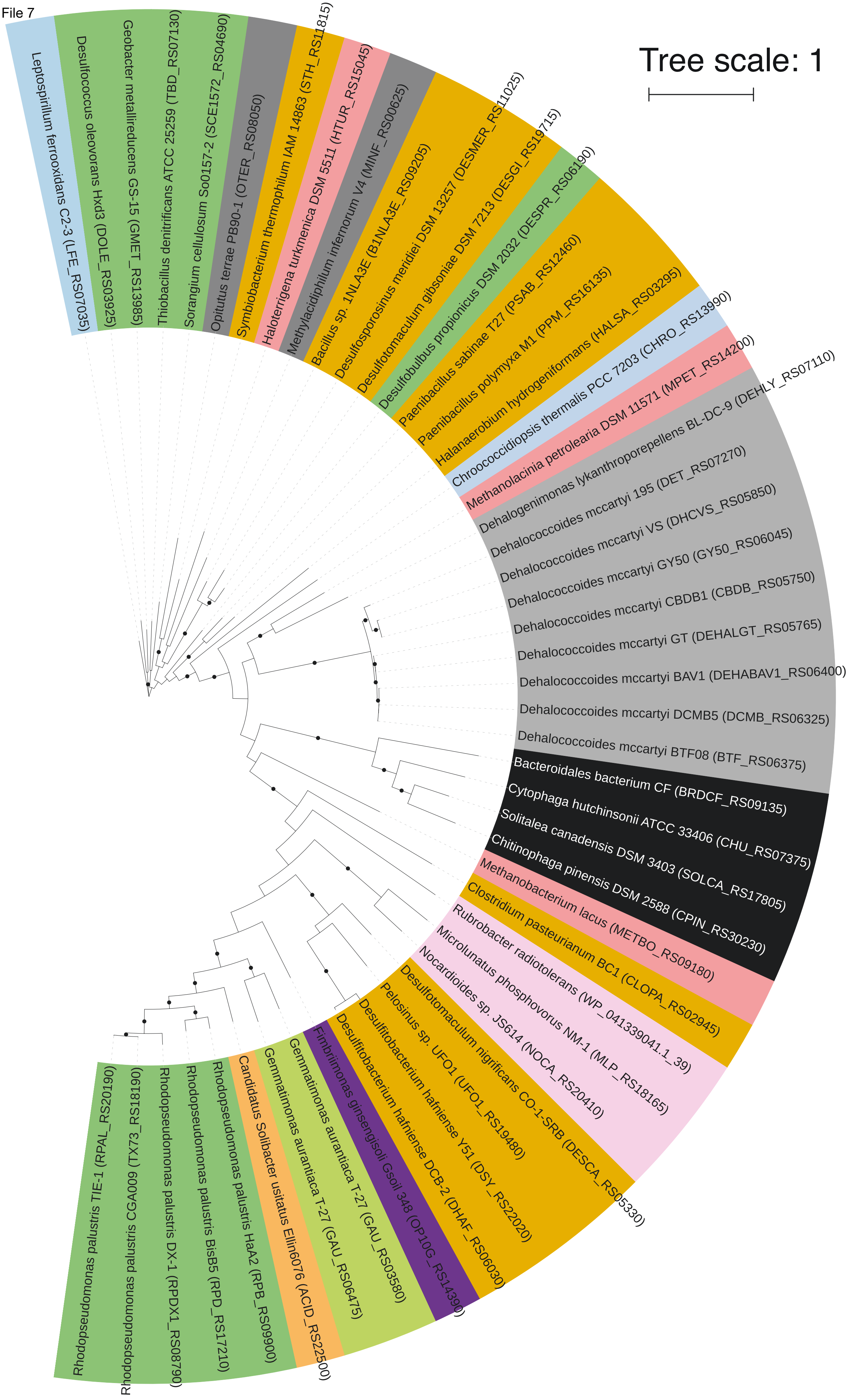

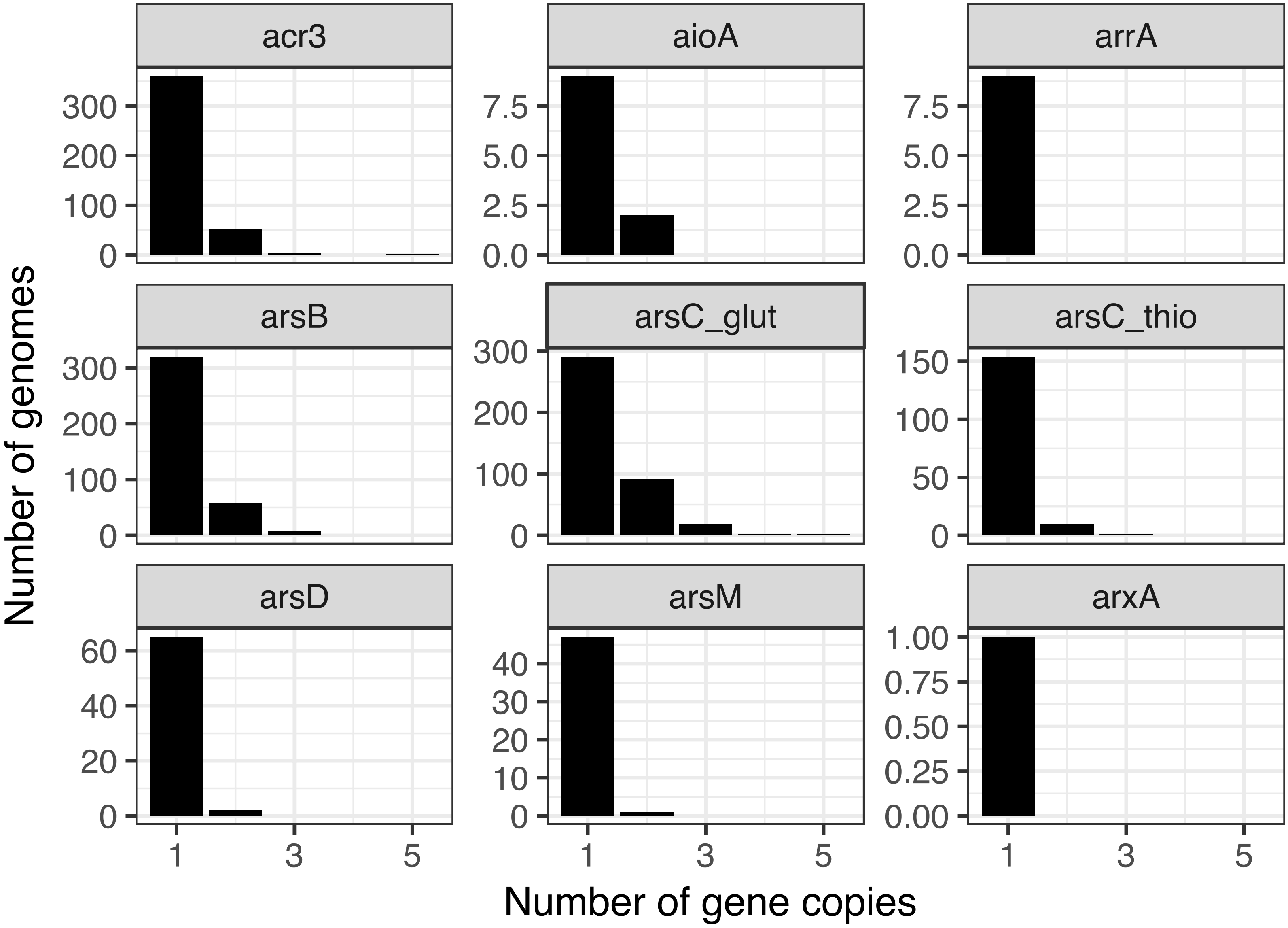
acr3
aioA
arrA
arsB
arsC_glut
arsC_thio
arsD
arsM
arxA
300
200
100
0
7.5
5.0
2.5
0.0
150
50
60
40
20
30
10
1.00
0.75
0.50
0.25
0.00
1
3
5
Number of genomes
Number of gene copies

Additional
File 9

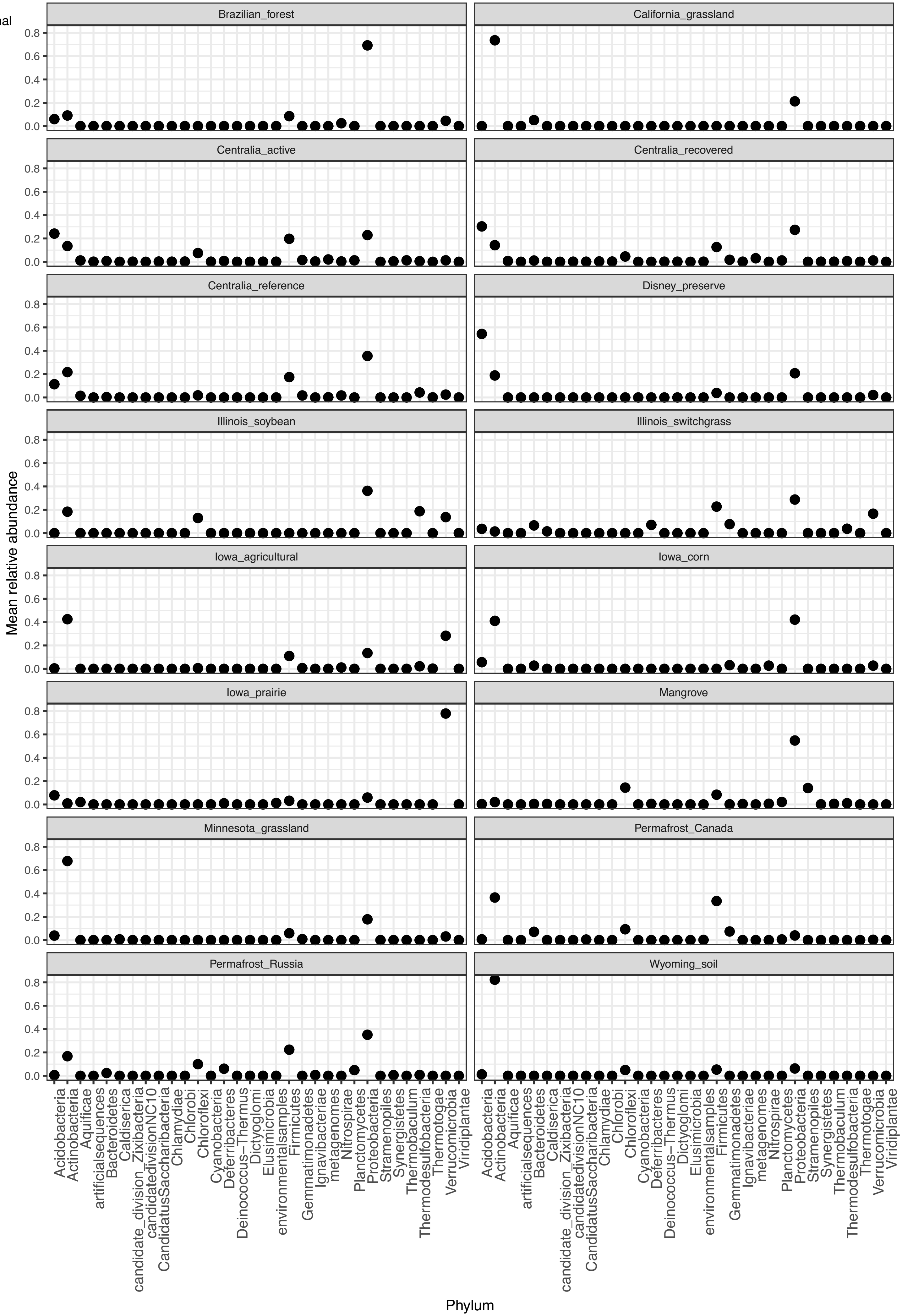

| Gene | Number of sequences | Number of endemic sequences | Percent endemic |
|---|---|---|---|
| *acr3* | 610 | 607 | 99.5 |
| *aioA* | 63 | 62 | 98.4 |
| *arrA* | 63 | 63 | 100 |
| *arsB* | 8 | 8 | 100 |
| *arsC* (grx) | 1316 | 1299 | 98.7 |
| *arsC* (trx) | 292 | 291 | 99.7 |
| *arsD* | 64 | 64 | 100 |
| *arsM* | 1193 | 1191 | 99.8 |
| *arxA* | 12 | 12 | 100 |
| **Totals** | 3621 | 3597 | 99.3 |

| FunGene Database/ Protein | Minimum HMM score | Minimum length (aa) | Minimum HMM coverage (%) | Number of FunGene sequences | Number of dereplicated sequences | Minimum assembled length (aa) |
|---|---|---|---|---|---|---|
| **ArsB** | 150 | 400 | 80 | 23680 | 5250 | 150 |
| **ACR3** | 140 | 300 | 80 | 19812 | 8002 | 150 |
| **ArsC_glut** | 80 | 120 | 85 | 18082 | 9635 | 50 |
| **ArsC_thio** | 172 | 100 | 80 | 7180 | 7180 | 50 |
| **ArrA** | 175 | 75 | 5 | 1621 | 1487 | 150 |
| **AioA** | 800 | 800 | 80 | 382 | 293 | 150 |
| **ArsM** | 200 | 100 | 30 | 3446 | 2948 | 160 |
| **ArsD** | 80 | 100 | 80 | 5404 | 876 | 150 |
| **ArxA** | 600 | 800 | 80 | 67 | 54 | 150 |
